# A Combined Chemo-Enzymatic Treatment for the Oxidation of Epoxy-Based Carbon Fiber-Reinforced Polymers (CFRPs)

**DOI:** 10.1101/2025.06.04.657587

**Authors:** Sasipa Wongwattanarat, Andrea Schorn, Leon Klose, Camille Carré, Ana Malvis Romero, Andreas Liese, Pablo Pérez-García, Wolfgang R. Streit

## Abstract

Carbon fiber-reinforced polymers (CFRPs), particularly epoxy-based composites, have become essential in the aerospace, automotive, and wind energy industries due to their robust mechanical properties, and lightweight nature. However, there is a lack of recycling technologies that are environmentally sustainable while also ensuring the recovery of carbon fibers in their original state. Although certain bacterial and fungal strains can colonize epoxy polymers, enzymes capable of efficiently degrading these materials have not yet been reported. Consequently, there is an urgent need for an effective, sustainable, and biologically inspired solution for CFRP recycling. Here, a chemo-enzymatic two-step oxidation process was developed. A chemical pre-treatment with propionic acid and hydrogen peroxide was used to recover imbedded carbon fibers. Additionally, three novel bacterial laccases isolated from the European spruce bark beetle (*Ips typographus*) demonstrated the ability to degrade three epoxy resin scaffolds from Hexflow® RTM6, used in aircraft applications. The sequential combination of both oxidative steps enabled the retrieval of clean carbon fibers and partial modification of epoxy functional groups, with the release of defined products over time. This bio-inspired approach renders the process more environmentally friendly and marks an initial step toward developing a bio-based recycling method for epoxy CFRPs.

## 1. Introduction

Epoxy polymers are thermosetting polymers that are essential in materials engineering due to their exceptional mechanical properties, as well as their high thermal and chemical resistance (Sukanto et al., 2021). These characteristics stem from a strong three-dimensional network formed during the curing process of epoxy resins and hardeners (Jin et al., 2015). Typically containing two or more oxirane groups, epoxy resins can be tailored to specific applications by combining them with various hardeners or curing agents like aliphatic and aromatic amines, anhydrides, thiols, and acids (Karak, 2021). This versatility makes them a preferred choice for carbon fiber-reinforced polymers (CFRPs) in industries such as aerospace, automotive, and construction, where lightweight, strength, and durability are crucial (Aamir et al., 2019; Ahmad et al., 2020). Notably, CFRPs constitute up to 53% by weight of current commercial aircraft structures (Gondaliya et al., 2016). As global demand for CFRPs rises by about 10% annually (Schüppel, 2023), with projected waste reaching 20 kt by 2025 (Zhang et al., 2020), effective management of their disposal has become a pressing concern.

Various recycling methods have been explored, with chemical recycling, or solvolysis, holding most promising for recovering both carbon fibers and matrix polymers (Liu et al., 2022). However, challenges remain in identifying environmentally friendly solvents that can operate under low temperature and pressure (Cheng et al., 2017). Despite progress, there is still a lack of sustainable, biologically inspired approach for degrading epoxy-based CFRPs (eCFRPs) (Klose et al., 2023).

The biodegradation of epoxy polymers remains challenging due to their synthetic nature: extensive cross-linking, stable chemical motifs (*i.e.* ethers and tertiary amines), and a lack of easily hydrolysable linkages. While some bacteria like *Pseudomonas* sp. and *Bacillus flexus* can colonize and reduce the corrosion resistance of epoxy coatings (Deng et al., 2019; Wang et al., 2016), significant advancements in identifying microorganisms capable of efficient degradation, similar to *Ideonella sakaiensis* with PET, have yet to be achieved (Yoshida et al., 2016). Though no enzyme has been identified to degrade epoxy, certain enzymes, particularly laccases (EC 1.10.3.2), have shown potential in partially and randomly oxidizing the surface of recalcitrant synthetic polymers (*e.g.*, PE, PVC) (Temporiti et al., 2022; Zhang et al., 2022), even though they do not significantly degrade the polymer backbone.

Laccases, or multi-copper oxidases (MCOs), can oxidize complex lignin-derived molecules with cross-linked ether and carbon-carbon bonds, as well as phenolic and non-phenolic compounds, by reducing molecular oxygen to water. These versatile enzymes catalyze various oxidative reactions, including phenolic oxidation, aromatic hydroxylation, oxidative polymerization, and demethylation (Bassanini et al., 2021; Munk et al., 2015), making them potential enzymes for modifying complex organic compounds with applications in biotechnology and environmental remediation (Khatami et al., 2022). Moreover, the laccase-mediator system (LMS) has evolved in nature to broaden their substrate range, with mediators (*e.g.*, ABTS (2,2’-azino-bis(3-ethylbenzothiazoline-6-sulfonic acid))) acting as electron shuttles (Hilgers et al., 2018). In contexts where natural enzymatic pathways struggle with recalcitrant synthetic polymers, laccases present a potential solution to the challenges posed by epoxy polymers.

This study developed a chemo-enzymatic treatment for the two-step oxidation of eCFRPs. Alternative pre-treatments were explored, with propionic acid/H_2_O_2_ at 65 °C and atmospheric pressure effectively breaking down diglycidyl ether of bisphenol F (DGEBF)-based composites while recovering carbon fibers close to their native state. Three novel bacterial laccases, isolated from the bark beetle (*Ips typographus*), were screened for their potential activity on three structural scaffolds of Hexflow® RTM6, a TGMDA-based epoxy commonly used in aerospace applications. Our findings indicate that the decomposition of epoxy via chemo-enzymatic treatment produces smaller compounds, representing an initial step toward developing a bio-based recycling strategy for eCFRPs.

## 2. Materials and methods

### 2.1 Bark beetle sample collection

Bark beetle specimens were collected using Theysohn slot traps (Niemeyer et al., 1983) baited with the beetle pheromone Ipsowit® (Witasek, Austria) and stored at -20 °C immediately after collection. The beetles were disinfected with 96% ethanol and affixed to paraffin plates with their ventral abdomen facing upward. During dissection, immobilized beetles were submerged in sterile 1X phosphate-buffered saline (PBS) pH 7.4. The intestines were extracted and stored at -20 °C.

### 2.2 DNA extraction and sequencing

Metagenomic sequencing of *Ips typographus* DNA was performed at the Heinrich Pette Institute in Hamburg, Germany. A genomic library was prepared using the NEBNext® Ultra™ DNA Library Prep Kit for Illumina® (New England BioLabs, Germany), and quality-checked with a BioAnalyzer High Sensitivity Chip (Agilent Technologies Inc., USA). Sequencing was performed on an Illumina HiSeq 2500 instrument (Illumina Inc., USA) in paired-end mode, generating 2 **×** 125 bp reads. Data were processed with Trimmomatic software v.0.32 (Bolger et al., 2014) for quality trimming and assembled using IDBA-UD software v.1.1.1 (Peng et al., 2012). The dataset was uploaded to the GOLD database (Kyrpides, 1999), annotated via the DOE-JGI pipeline (Huntemann et al., 2015), and stored in the IMG database (Markowitz et al., 2012) with gene ID: ItL-01, Ga0063521_10002204; ItL-02, Ga0063521_100014138; ItL-03, Ga0063521_100024328; ItL-04, Ga0063521_1000001282; ItL-05, Ga0063521_100021622; ItL-06, Ga0063521_100028714. A search was performed in the IMG metagenome using the keyword “multicopper oxidase”. Six candidate genes, with completeness and the presence of all four conserved copper-binding site (CBS) motifs, are listed in Table S1.

### 2.3 Bioinformatic analysis

Amino acids of the putative and the recognized laccases were acquired from public sequence databases NCBI, UniProt, and IMG (”Database resources of the National Center for Biotechnology Information,” 2018; Markowitz et al., 2012; The UniProt Consortium, 2017). Local alignments were performed with BLASTp (Boratyn et al., 2012). The sequence alignment of the amino acid sequences was conducted using the T-Coffee server with Expresso mode (Armougom et al., 2006). The phylogenetic tree was constructed using MEGA11 with maximum-likelihood method and JTT matrix-based model with 1,000 bootstrap replicates (Tamura et al., 2021). The 3D structural models were predicted using AlphaFold 3 (Abramson et al., 2024). The evolutional conservation profiles of the proteins were analyzed using Consuf server (Yariv et al., 2023). The models were visualized using UCFS Chimera v.1.16 (Huang et al., 2014).

### 2.4 Molecular cloning, protein expression, and purification

The MCO genes were synthesized with codon optimization for *E. coli* (MWG Eurofins, Germany). The strains, plasmids and primers used are listed in Table S2. Genes were inserted into pET21a(+) expression vectors. ItL-01-03 were transformed into *E. coli* BL21(DE3) (Novagen/Merck, Germany), while CueO was transformed into *E. coli* T7 Shuffle (New England BioLabs, Germany). An overnight inoculum (1%) was grown aerobically in autoinduction medium (ZYM-5052; Studier (2005)), with 100 μg/mL ampicillin at 37 °C until an OD_600_ of 0.6 was reached. Cultures were then supplemented with 250 µM CuSO_4_ and incubated at 28 °C for 16 to 20 hours. Cells were harvested, treated with 1 mM phenylmethanesulfonyl fluoride (PMSF), and lysed three times at 1,250 psi using a French press (American instrument, USA). Proteins were purified with Ni-NTA agarose (Qiagen, Germany) and dialyzed in 50 mM Tris-HCl pH 7 using a 30 kDa Amicon tube (GE Health Care, Germany).

### 2.5 Biochemical characterization

A series of spectrophotometric assays using 2,2′-azino-di-(3-ethylbenzthiazoline sulfonic acid) (ABTS; Merck, Germany) was performed to characterize the recombinant laccases, with a detectable color at 420 nm (ε = 36,000 M^-1^ cm^-1^), measured using a Synergy HT microplate reader (BioTek, Germany).

In each assay, 2 µM laccase was incubated with 1 mM ABTS in a total volume of 200 µL within a 96-well plate, supplemented with 1 mM CuSO_4_. Initial tests determined optimal pH using 0.1 M citrate-phosphate buffers from pH 3 to 7, followed by temperature optimization from 20 to 80 °C. Buffer preference was evaluated with 0.1 M citrate-phosphate and acetate buffers at their respective optimal pH levels. Thermostability assays were conducted at 30 to 60 °C for seven days. The effect of Cu^2+^ supplementation (0 µM to 1 mM) was tested in 0.1 M acetate buffer at pH 4 and optimal temperatures (T*_opt_*).

Kinetic constants were determined using ABTS in 200 µL, 0.1 M acetate buffer pH 4 at 25 °C for 30 minutes. The Michaelis-Menten constant (*K*_m_), maximum reaction rate (*V*_max_), and turnover rate (*K*_cat_) were calculated by fitting the initial rates to the Michaelis-Menten equation with Solver (Microsoft Excel add-in, Frontline Systems, Inc., USA) (Fig. S3) (Chris & Nithesh Chandrasekharan, 2020).

Laccase ItL-03 was characterized for optimal pH with substrates ABTS, syringol, guaiacol (Merck, Germany) using 0.1 M buffers: citrate-phosphate (pH 3-6), phosphate (pH 6-8), Tris (pH 8-9), and carbonate-bicarbonate (pH 9-10), with 3 mM substrate and 5 mM CuSO_4_. Specific activity (U/mg) was evaluated with 1 mM substrate and 1 mM CuSO_4_, monitoring absorbance over 3 minutes at ε420 = 36,000 M^-1^ cm^-1^ for ABTS, ε468 = 14,800 M^-1^ cm^-1^ for syringol, and ε465 = 12,000 M^-1^ cm^-1^ for guaiacol.

### 2.6 Assessment of enzymatic activities on epoxy surrogates

The oxidative activity of laccases on bis(4-dimethylamino-cyclohexyl) methane (BBCM; kindly donated by the Institute of Technical Biocatalysis, Hamburg University of Technology, Germany), 1,3-bis(methyl(phenyl)amino) propan-2-ol (BMAP; kindly contributed by the Manchester Institute of Biotechnology, University of Manchester, UK), and N, N-bis(2-hydroxypropyl)-p-toluidine (NNBT; Toronto Research Chemicals, Canada) was investigated. These substrates were dissolved in dimethyl sulfoxide (DMSO) to prepare 100 mM stock solutions. In each reaction, 0.02 U/mg of purified enzyme was incubated with 3 mM substrate, 1 mM CuSO_4_, and 1 mM ABTS in 0.1 M acetate buffer pH 4 at T*_opt_*. The mixture was shaken at 450 rpm for 2 hours. For endpoint analysis, samples were diluted 1:3 with buffer, centrifuged at 13,000 rpm for 6 minutes, then extracted with dichloromethane (DCM) in a 1:1 ratio and centrifuged again. The DCM layer was diluted 100-fold in LC-MS grade water.

#### 2.6.1 Liquid chromatography-mass spectrometry (LC-MS)

LC analysis was performed using a Dionex Ultimate 3000 UHPLC system with an Agilent Zorbax Extend-C18 column (2.1 **×** 50 mm, 1.8 µm). The mobile phase was acetonitrile and water with 0.1% (v/v) formic acid, with a gradient from 5% to 95% acetonitrile over 28 minutes at a flow rate of 0.3 mL/min, monitored at 254 nm. Mass detection used a Bruker maXis ESI-QTOF in positive mode (*m/z* 50-2300, capillary voltage 4 kV). Data were analyzed using MestReNova x64 (Mestrelab Research S.L.U, Spain). For assessing ItL-03 degradation of NNBT, LC system was replaced with an Agilent 1260 HPLC, and the run time was extended to 30 minutes, with other parameters remained unchanged. Analyte concentrations were determined via calibration curves (Fig. S5). The initial concentration at t_0_ was set at 100%, and the degradation rate was calculated by subtracting the remaining epoxy from 100%.

### 2.7 Pre-treatment analysis of CFRPs

Carbon fiber-epoxy resin (eCFRPs) was purchased from Goodfellow, Ltd., Germany (product number C-42-SH-000150), comprising Toray T300 carbon fiber (or equivalent) with 50% volume fraction and Elantas EC157 epoxy resin (or equivalent). The diglycidyl ether of bisphenol F (DGEBF)-based composite, 0.5 mm thickness and 150 **×** 150 mm, was laser-cut into 6 **×** 12 mm pieces. Samples were incubated in various acidic-peroxide solutions, containing either 5 M or 9 M acid and H_2_O_2_ in a 95:5 ratio. The solution volume was 60 mL/g, and treatments lasted 8, 24, and 48 hours at 65 °C with continuous shaking at 200 rpm. Control samples were exposed to acid, H_2_O_2_, and water. After treatment, samples were washed thoroughly with warm water and dried overnight at 60 °C. The mass fraction of resin was determined following DIN EN 2564:2018 (e.V., 2019). The resin weight loss rate was calculated based on established methods (Das et al., 2018).

#### 2.7.1 Fourier transform infrared (FTIR) spectroscopy

FTIR spectra were recorded using a Vertex 70v spectrometer (Bruker, Germany) in ATR mode. Pre-treatment analysis of CFRPs was analyzed over a spectral range of 4,000 to 650 cm^-1^, at a resolution of 2 cm^-1^, with each spectrum generated from 50 scans at 27 °C. The measurements of the RTM6 powder used in chemo-enzymatic treatment were conducted similarly, but over a range of 4,000 to 600 cm^-1^ with 32 scans. Data were processed using OPUS software (Bruker, USA), normalized to [0, 1], and smoothed with a Savitzky-Golay filter (20 points) using OriginPro 2024 (OriginLab Inc., USA).

#### 2.7.2 Scanning electron microscopy (SEM) analysis

Imaging was performed with a LEO 1525 Field Emission Scanning Electron Microscope (LEO Electron Microscopy Inc., USA), operating at an electron high tension (EHT) of 5-10 kV and a working distance (WD) of 8.1-9.6 mm. Data was processed using SmartSEM V06.00 software (Carl Zeiss Microscopy GmbH, Germany).

#### 2.7.3 Energy-dispersive X-ray (EDX) spectroscopy

Elemental analysis of the carbon fibers recovered from pre-treatment was carried out with the Oxford XMax 50 EDX measuring device from Oxford Instruments in combination with a scanning electron microscope (SEM, Auriga 40 from Zeiss).

### 2.8 Chemo-enzymatic oxidation

HexFlow®-like RTM6 cured resin was synthesized and ground into a powder (particle size ≤ 0.6 mm). The tetraglycidyl methylene dianiline (TGMDA)-based powder was immersed in 5 M propionic acid/H_2_O_2_ (95:5) at 60 mL/g, incubated for 48 hours at 65 °C with shaking at 230 rpm. The resulting solutions were neutralized with 5 M NaOH and extracted with DCM (1:1). After evaporating the DCM, the precipitate was resuspended in DMSO at a DCM:DMSO ratio of 10:1 based on the original DCM volume. This is referred to extracted epoxy. The remaining powder was filtered, washed with water, and dried overnight at 60 °C.

10 µL of extracted epoxy was combined with 0.1 U/mg ItL-03, 5 mM CuSO_4_, and 5 mM ABTS or no mediator in 0.1 M acetate buffer at pH 4, shaken at 900 rpm at 50 °C. Samples were collected after 2 and 24 hours, with bovine serum albumin (BSA) serving as a control. Samples were extracted with DCM (1:1), centrifuged at 13,000 rpm for 5 minutes, and the organic phase collected for MS analyses.

15 mg of non-treated and pre-treated epoxy powder was incubated with 0.05 U/mg ItL-03 for 5 days under the same settings. Mediators, including ABTS, syringol and (2,2,6,6-tetramethylpiperidin-1-yl)oxidanyl (TEMPO; Thermo Fisher Scientific, Germany), were tested at their optimal pH (Fig. S8), with TEMPO at pH 4 (Azimi et al., 2016; Wong et al., 2013). After enzymatic treatment, powder was washed with water and dried overnight at 60 °C for FTIR analysis.

#### 2.8.1 Direct electrospray ionization-mass spectrometry (ESI-MS)

ESI-MS analysis was performed via direct injection on an Agilent 6224 ESI-TOF coupled with an Agilent 1200 HPLC. Ionization in positive mode used a 4 kV spray voltage, with a scan range of *m/z* 110 to 3,200. Flow rate was 0.3 mL/min, with autosampler cooled to 15 °C. Drying gas flow was 10 L/min, nebulizer pressure at 15 psi, and gas temperature at 325 °C. Data were analyzed with MestReNova x64 software (Mestrelab Research S.L.U, Spain).

#### 2.8.2 Gas chromatography -mass spectrometry (GC-MS)

GC-MS analysis was performed on an Agilent GC 7890A GC coupled with an Agilent 5975C VL-MSD. A Thermo Fisher Scientific TG-5MS column (30 m **×** 0.25 mm, 0.25 µm) was used. The program started at 80 °C for 1 minute, then ramped at 10 °C/min to 300 °C, and held for 10 minutes. Inlet temperature was 250 °C; helium (5.0) at 1 mL/min with a 1:10 split ratio. Data were acquired in full scan mode (*m/z* 35 to 500) with the ion source at 230 °C, and processed using MestReNova x64 software (Mestrelab Research S.L.U, Spain).

## 3. Results

### 3.1 Bark beetle metagenome uncovers novel bacterial laccases

European spruce bark beetles colonize and feed on the inner bark layer of trees, a behavior typical of xylophagous beetles that exhibit laccase-like activity in their digestive systems (Christiansen & Bakke, 1988; Geib et al., 2008). This species represents a promising source of enzymes with lignolytic properties, such as laccases.

The bark beetles were collected in a forest near Hanover, Germany, using pheromone traps, and their gastrointestinal tracts were isolated for microbiota DNA extraction and sequencing (Fig. 1a). The assembled metagenomic data, approximately 219 Mbp with a total of 434,200 protein coding genes, were searched for multicopper oxidases (MCOs) or laccases, yielding six putative MCOs designated ItL-01 to ItL-06. However, only ItL-01 to ItL-03 were successfully expressed and produced (Fig. S1).

**Figure 1.**
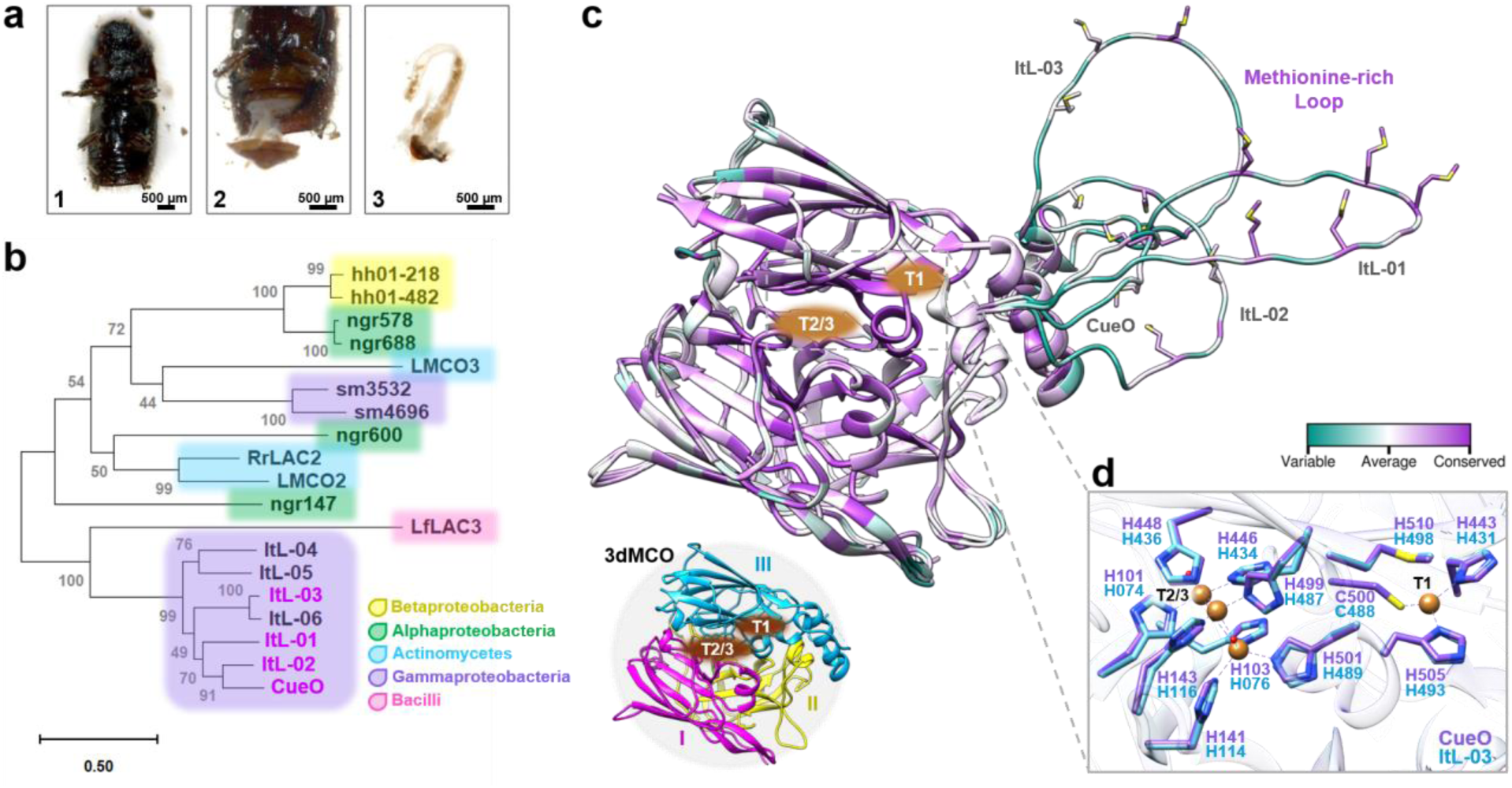
Putative bacterial laccases identified in the gut metagenome of the bark beetle *Ips typographus*. (a) Dissection of the bark beetle intestines. The abdomen (1) was opened with sterilized tweezers, (2) and the intestines were removed, (3) and collected in sterile phosphate-buffered saline for DNA extraction. (b) A phylogenetic tree of putative MCOs based on amino acid sequences from beetles and bacteria associated with lignin and synthetic polymer degradation. The accession numbers, UniProt or IMG entries for the sequences used are listed in Table S1. (c) The alignment of the 3D structures of CueO and ItL-01-03 shows the conservation of amino acid positions, represented as B-factors. Structural details highlight the copper binding sites (CBSs) T1 and T2/3 sites, and the methionine-rich loop. These MCOs, represented by CueO (PDB: 4NER), consist of three cupredoxin-like domains (3dMCO), labelled I, II, and III. (d) Due to the similarities in CBSs between CueO and ItL-01-03, only CueO (purple) and ItL-03 (blue) are shown.

A phylogenetic tree was constructed based on protein sequences to assess the evolutionary relationships among these laccases, the benchmark laccase CueO from *E. coli* (Blattner et al., 1997; Grass & Rensing, 2001), eight recognized laccases associated with lignin-rich environments or plant pathogenesis (Avison et al., 2000; Hornung et al., 2013; Trinick, 1980), and three laccases reported to partially degrade polyethylene (PE) (Fig. 1b) (Zampolli et al., 2023; Y. Zhang et al., 2023). Phylogenetic analysis revealed that these MCOs are predominantly affiliated with the phylum Pseudomonadota, with few belongings to Actinomycetota and Bacillota, aligning with the taxonomic classification of bacterial laccases in the LccED database (Gräff et al., 2020). ItL-01 to ItL-06 exhibit over 60% sequence similarity to CueO, while the other laccases have identities below 40% (Table S1).

The 3D structural analysis indicates that these laccases are typical three-domain MCOs (3dMCO), with highly conserved regions near the two copper binding sites (CBSs) at the T1 and T2/T3 centers (Fig. 1c). The T1 copper ion is located in the third domain, where the substrate is oxidized, while the T2 and T3 copper ions interface between the first and third domains, where dioxygen reduction takes place (Gräff et al., 2020). In contrast, methionine-rich loops near the T1 site, are considerably variable. In fact, the methionine residues themselves are quite conserved. These methionines are thought to facilitate the recruitment and transport of copper (Contaldo et al., 2024).

Crystals of CueO (PDB 4NER; Komori et al. (2014)), with a resolution of 1.60 Å and containing copper (II) ions, were used to map CBSs with ItL-03, showing high similarity and conservation among MCOs, predominantly with histidine residues (His) (Fig. 1d). Given the highly conserved topology of CBSs, the methionine-rich loops may influence enzyme specificity and reactivity, *e.g*., regulating substrate access and stabilizing substrates (Borges et al., 2020).

### 3.2 Biochemical characterization identifies thermostable laccases functioning at low pH

Recombinant enzymes CueO and ItL-01-03 were produced and purified via affinity chromatography, yielding predicted molecular weights of 50-60 kDa (Fig. S1). ABTS (2,2′-azinobis (3-ethylbenzothiazoline-6-sulfonic acid)), a common mediator in laccase/mediator systems and a typical substrate for laccases (Hilgers et al., 2018), was used to measure radical formation at 420 nm to investigate enzyme characteristics.

The bacterial laccases showed a preference for the acidic pH range, maintaining over 50% activity between pH 3.0 and 4.6 (Fig. 2a). ItL-01 reached a pH maximum of 4.2, while CueO, ItL-02, and ItL-03 peaked at pH 3.8. Particularly, ItL-03 retained over 90% activity between pH 3.0 and 3.8, indicating pronounced acidophilic nature. Since the optimal pH range for these enzymes was similar, acetate buffer at pH 4.0 was selected for further characterization (Fig. S2).

**Figure 2.**
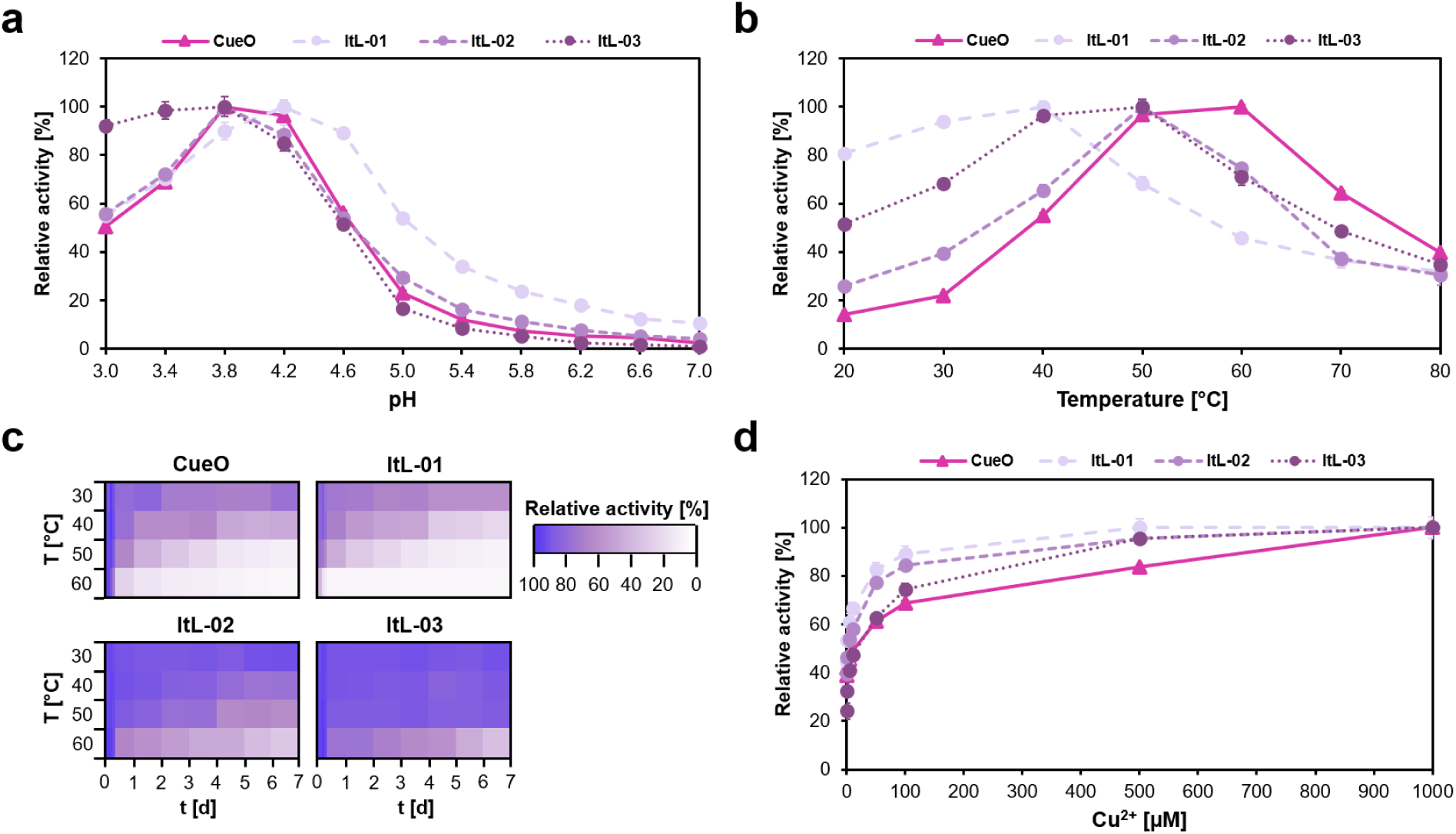
Biochemical characteristics of CueO and ItL-01-03, determined using ABTS, highlight their stability and limitations in synergistic enzyme activity. (a) pH profile: the enzymes preferred low pH and reached their maximum activity at pH 4.0. (b) Temperature profile: each enzyme exhibited a distinct temperature stability, within the mesophilic range of 30-60 °C. (c) Thermostability: most enzymes retained their activity for only a few hours, losing nearly 80% within a day at elevated temperatures, while ItL-02-03 maintained over 60% activity at 50 °C for a week. (d) Effect of copper ions (Cu^2+^): the activity of laccases significantly increases with higher Cu^2+^ when supplemented during enzymatic reactions. Error bars indicate the standard deviation (*n* = 3). The standard deviation in ‘c’ was below 6%. Buffer preference is shown in Fig. S2.

Temperature profiles and thermostability were assessed by incubating enzymes at 30-60 °C for 7 days. The bacterial laccases exhibited moderate optimal temperatures (T*_opt_*) (Fig. 2b). ItL-01 had a T*_opt_* of 40 °C, retaining over 80% activity after 10 hours (Fig. 2c). CueO exhibited the highest T*_opt_* at 60 °C, but experienced a 30% drop-in activity after 10 hours and nearly lost all activity after three days. ItL-02 and ItL-03 shared the same T*_opt_* of 50 °C. While ItL-02 retained 80% activity at its T*_opt_* after two days, ItL-03 sustained this level even after seven days (Fig. 2c), making it the most thermophilic and thermostable enzyme in this study.

To ensure proper folding, 0.25 µM CuSO4 was supplied during enzyme production. Supplementing additional Cu^2+^ into the reactions resulted in a gradual increase in laccase activity (Fig. 2d). The steady-state kinetics of the laccases were determined with respect to ABTS, revealing similar catalytic turnover values (Table S3). Among all, ItL-03 exhibited the highest catalytic turnover (*k*_cat_ = 3.04 min^-1^) but had relatively high *K*_m_ value of 7.76 mM. However, in terms of catalytic efficiency (*K*_cat_/*K*_m_), ItL-03 outperformed the other laccases with a value of 0.39 min^-1^ mM^-1^. Based on these initial characterizations, the laccases were further evaluated for their catalytic properties toward epoxy model building blocks.

### 3.3 Oxidoreductases can degrade epoxy resin building blocks

The laccases were tested for their ability to degrade bis(4-dimethylamino-cyclohexyl) methane (BBCM), 1,3-bis(methyl(phenyl)amino) propan-2-ol (BMAP), and N, N-bis(2-hydroxypropyl)-p-toluidine (NNBT) (Fig. 3). These substrates contain key resin motifs, including a tertiary amine found in HexFlow® RTM6, a widely used aerospace epoxy resin (Zotti et al., 2020), serving as models to study enzyme mechanisms on C-N bonds in the original polymer.

**Figure 3.**
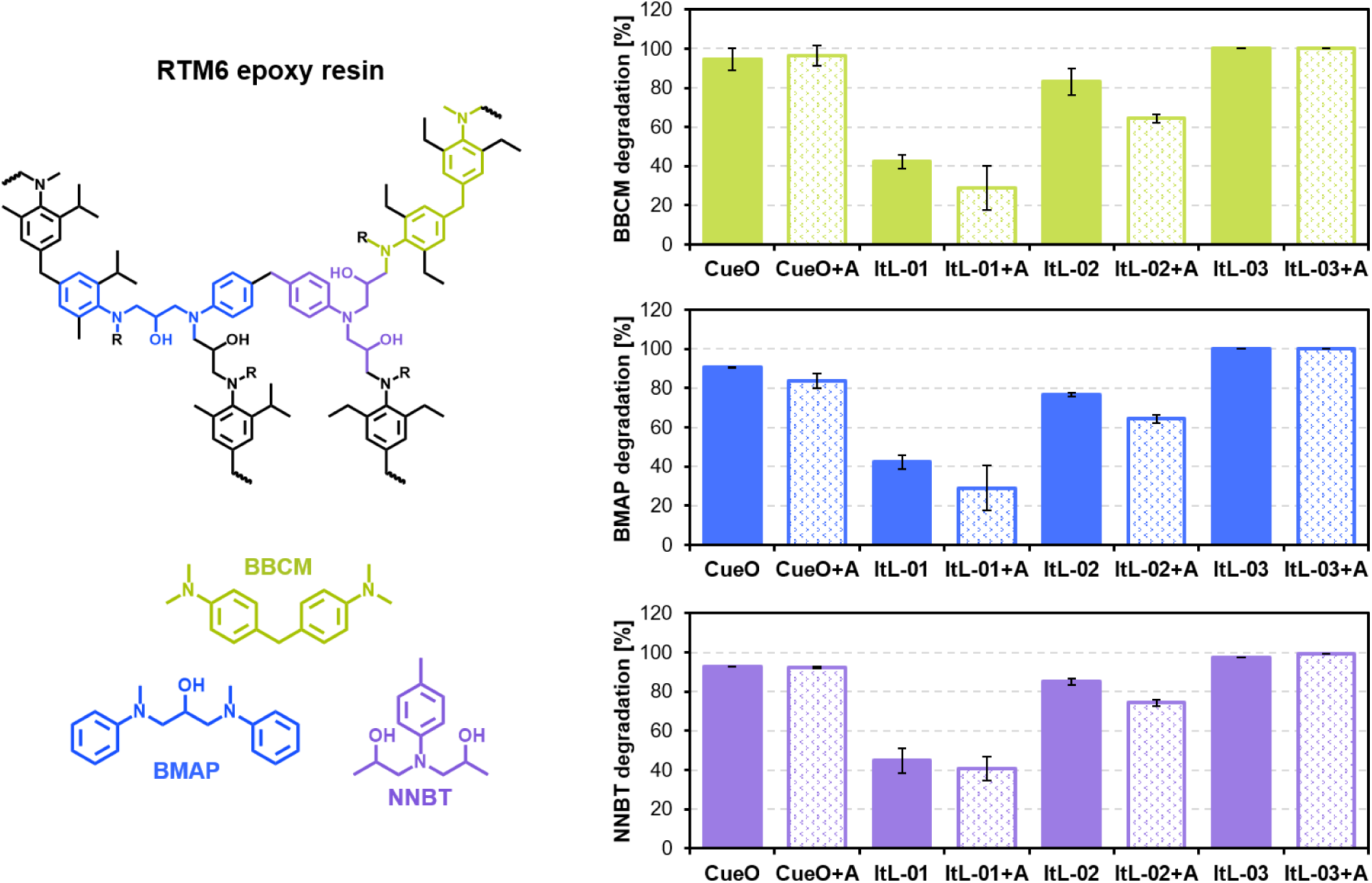
Bacterial laccases degraded the epoxy scaffolds—BBCM, BMAP, and NNBT—derived from Hexflow® RTM6 epoxy resin. The tests were conducted in the presence and absence of 1 mM ABTS (A) at their respective optimal pH and temperature conditions, and the remaining concentration of the epoxy substrate after 2 hours was measured using LC-MS (Fig. S4). The initial observation (t_0_) defined the starting amount of epoxy, set at 100%. The degradation rate was calculated by subtracting the percentage of remaining epoxy from 100%. Error bars indicate the standard deviation (*n* = 3). Supporting data can be found in Fig. S4-7.

Degradation of these epoxy scaffolds was monitored after 2 hours using liquid chromatography-mass spectrometry (LC-MS) (Fig. S4). The laccases exhibited comparable activity levels across the different substrates. ItL-03 showed the highest activity, completely converting all three substrates BBCM, BMAP, and NNBT within 2 hours (Fig. 3), and achieving 80% conversion after just 30 minutes (Fig. S6). This was followed closely by CueO, ItL-02, and ItL-01, with conversions of approximately 90%, 80%, and 40%, respectively. LC-MS analysis indicated possible N-dealkylation activity of the enzymes (Fig. S7).

Given that laccase activity could be enhanced by a mediator, the effect of ABTS (A) were evaluated. The presence of ABTS only slightly affected the enzymes’ activity, resulting in either a marginal decrease or nearly the same level of activity (Fig. 3, +A). This is likely because the small model substrates can access the enzyme’s active site directly, making a mediator unnecessary. Among the bacterial laccase candidates, ItL-03 emerged as the most effective for degrading the epoxy scaffolds, prompting further analysis of its efficacy on cured epoxy.

The activity of laccase ItL-03 was further investigated with additional mediators, syringol and guaiacol. The optimal pH and buffer preferences for each mediator were determined (Fig. S8), and the specific activities of ItL-03 with ABTS, syringol, and guaiacol were assessed (Fig. S9a). ItL-03/syringol exhibited the highest activity, approximately four times greater than that of ABTS, while guaiacol showed relatively low specificity. The optimal concentration of copper was further determined to be 5 mM for ItL-03 activity (Fig. S9b).

The activity of ItL-03 on NNBT with mediators was evaluated (Fig. S10). ItL-03/guaiacol oxidized NNBT more rapidly than with ABTS, achieving 90% conversion in 30 minutes, although similar conversion levels were reached after two hours. Since guaiacol is a phenolic mediator similar to syringol but with a slightly lower oxidation potential, and achieves comparable NNBT conversion levels to ABTS, it was excluded from further investigation into epoxy degradation.

Initial assays on eCFRPs, however, did not result in any observable degradation. Therefore, pre-treatment was investigated to determine its potential to enhance enzyme access to resulting oligomers or degradation intermediates.

### 3.4 Propionic acid-peroxide recovers clean carbon fibers and depolymerizes epoxy based CFRPs

Conventionally, the European standard (DIN EN 2564:2018) employs concentrated sulfuric acid and 30% hydrogen peroxide (H_2_O_2_) to treat CFRPs at high temperatures (160-260 °C) to determine fiber, resin, and void contents. Due to high corrosiveness and toxicity, alternative organic acids— formic acid (FA), acetic acid (AA), propionic acid (PA), lactic acid (LA), malic acid (MA), tartaric acid (TA), citric acid (CA)—combined with H_2_O_2_ were selected for eCFRP pre-treatment. These acids are less toxic, derived from renewable sources, and biodegradable.

Diglycidyl ether of bisphenol F (DGEBF)-based composites were treated at 60 °C for 24 and 48 hours in a 5 M acidic-peroxide mixture with a 95:5 ratio. This specific ratio minimizes damage to carbon fibers due to the limited amount of H_2_O_2_ (Das et al., 2018). Their remaining weights, after pre-treatment, were calculated to evaluate the efficacy of different acids. All treatments with H_2_O_2_, except for NA, exhibited higher resin decomposition than those without (Fig. 4a). This highlights the importance of H_2_O_2_ in the reaction mechanism, probably due to the *in-situ* formation of peracids. AA- and PA-H_2_O_2_ achieved 40% and 80% epoxy resin degradation after 24 hours, respectively, with the latter capable of complete degradation after 48 hours. Using 9 M AA- and PA-H_2_O_2_ enabled effective depolymerization within 8 hours (Fig. S11). Conversely, high molecular weight organic acids (LA, MA, TA, CA) and SA showed minimal oxidative activity towards the composites (Fig. 4a).

**Figure 4.**
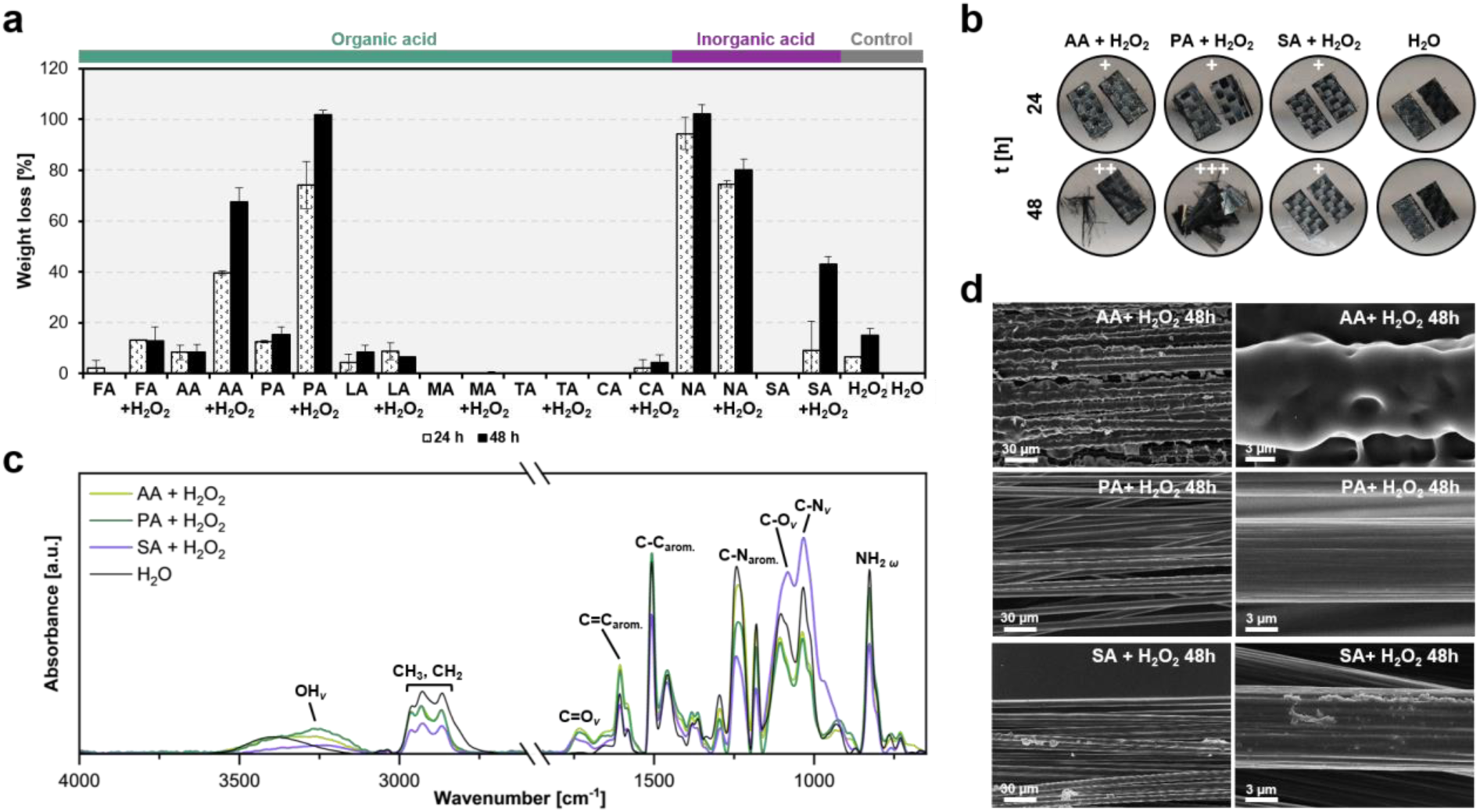
Organic acid treatments are effective in degrading epoxy polymers and recovering clean carbon fibers. (a) Weight loss of the resin mass in eCFRPs treated with organic acids, with and without hydrogen peroxide (H_2_O_2_), was compared to inorganic acids and controls (H_2_O and H_2_O_2_). The acids tested included formic acid (FA), acetic acid (AA), propionic acid (PA), lactic acid (LA), malic acid (MA), tartaric acid (TA), citric acid (CA), nitric acid (NA), and sulfuric acid (SA). Composites were incubated in a 95:5 acid-to-peroxide ratio at a 5 M, 65 °C, and 200 rpm for 24 and 48 hours. Error bars indicate the standard deviation (*n* = 3). (b) Morphological changes in the composites after 24 and 48 hours of pre-treatment. +, ++, and +++ represent the degree of carbon fiber exposure, ranging from low to high. (c) FTIR analysis of the 8-hour pre-treatment of eCFRPs. The corresponding functional groups are indicated. *v* denotes stretching, while *ω* denotes wagging. (d) SEM images of the eCFRPs after 48-hour treatments. Scale bars indicate size in micrometers (µm). Supporting data can be found in Fig. S11-14.

Carbon fibers (CFs) became visible in composites treated with PA-H_2_O_2_ after 48 hours (Fig. 4b and S12). Scanning electron microscopy (SEM) images revealed clean and elongated CFs from the PA-H_2_O_2_ treatment comparable to those treated under the European protocol (Fig. 4d and S13). Energy Dispersive X-ray (EDX) analysis confirmed that CFs recovered after PA-H_2_O_2_ predominantly contained carbon (C), with negligible oxygen content likely due to manufacturing impurities (Fig. S14).

Fourier Transform Infrared Spectroscopy (FTIR) analysis was performed to examine changes in functional groups following pre-treatments. The C=C aromatic band (around 1,600 cm^-1^) was evident in all epoxy composites, while the C=O stretching band (around 1,750 cm^-1^) appeared only in samples subjected to acid-peroxide pre-treatment, indicating possible oxidation (Fig. 4c). AA- and PA-H_2_O_2_ exhibited higher peak intensities in the C=C region than SA-H_2_O_2_, possibly due to the formation of carbonyls. Epoxy functional groups (*e.g*., C–N stretching in primary aliphatic and aromatic amines, around 1,030 cm^-1^ and 1,240 cm^-1^ respectively) significantly decreased in the AA- and PA-H_2_O_2_ treated samples (Fig. 4c). The stretching peaks of OH and C=O bonds in epoxy treated with organic acid-H_2_O_2_ were more pronounced than those in SA-H_2_O_2_, which was more effective at converting C–N in aromatic amines to C–N in primary amines. This suggests that organic and inorganic acids target different functional groups.

These findings demonstrate that PA-H_2_O_2_ pre-treatments effectively facilitate epoxy degradation, that combines effectiveness to lower environmental risks. This pre-treatment allows for the recovery of clean CFs, which could potentially be reused.

### 3.5 A two-step oxidative process transforms RTM6 epoxy into defined products

The PA-H_2_O_2_ pre-treatment effectively depolymerized the epoxy matrix of the DGEBF-based composite and recovered clean CFs. While CFs can be retrieved using the pre-treatment, it is important to determine whether laccase ItL-03 can further modify the remaining polymeric matrix to yield a more defined educt for downstream processing.

To evaluate the potential for enhanced bio-based recycling through chemo-enzymatic oxidation, Hexflow®-like RTM6 powder was used, synthesized from TGMDA amine epoxy precursor and the di-amine hardeners 4,4′-methylenebis(2,6-diethylaniline) (MDEA) and 4,4′-methylenebis(2-isopropyl-6-methylaniline) (M-MIPA) (Addou et al., 2016). RTM6 epoxy was selected over the composite, as the CFs were no longer considered, and it also enabled the investigation on this high-performance amine epoxy. Following PA-H_2_O_2_ pre-treatment of the epoxy powder, the resulting solution was neutralized and extracted, referred to as “extracted epoxy”. This extracted epoxy was subsequently introduced to ItL-03, and the resulting samples were analyzed using electrospray ionization-mass spectrometry (ESI-MS) via direct injection and gas chromatography-mass spectrometry (GC-MS).

ESI mass spectra revealed that the PA-H_2_O_2_ treatment effectively depolymerized epoxy resins into several small compounds within mass-to-charge ratio (*m/z*) ranges of 150-200 and 300-500, (Fig. S15). However, after 24 hours of treatment with ItL-03, the ions at *m/z* 483.3 were present but at a relatively low abundance, especially with the addition of the mediator ABTS, when compared to the BSA and buffer controls (t_0_ and t_24_) (Fig. 5a). Additionally, ions at *m/z* 354.2 and 501.3 were detected at lower intensities in the laccase samples, regardless of ABTS presence (Fig. 5b and S16). The reduced abundance of ions at *m/z* 354.2 and 483.3 in the presence of laccase ItL-03 suggests potential degradation or oxidation by the enzyme. Fig. 5c proposes potential structures, considering adduct formation.

**Figure 5.**
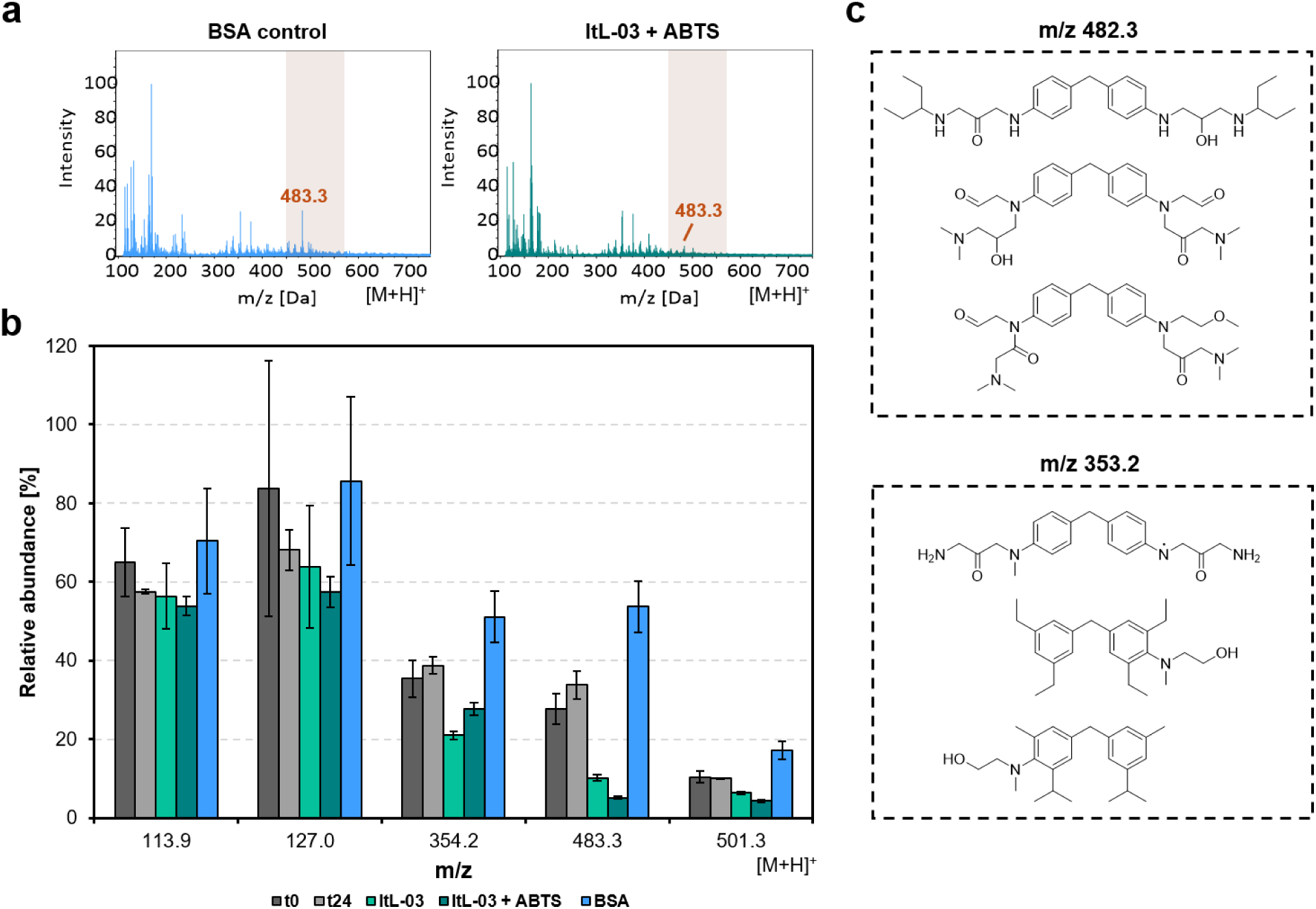
ESI-MS analysis of extracted epoxy from chemo-enzymatic degradation using PA-H_2_O_2_ and laccase ItL-03 on RTM6. (a) ESI-MS spectrum obtained via direct injection in positive mode, showing the degradation activity of ItL-03 with ABTS and BSA control after 24 hours. *m/z* 483.3 was mostly absent in the laccase sample but remained present in the control. (b) The relative abundance of *m/z* values of interest from the same experiment in ‘a’ is shown, including other samples. The spectra were normalized using the *m/z* value at 163.9 as a reference peak. t_0_ and t_24_ represent controls without enzyme at the beginning and end of the incubation period. Error bars represent the standard deviation (*n* = 3). (c) Proposed compounds may be degraded by the enzyme, with the *m/z* values of 482.3 and 353.2, after considering adduct formation. Supporting data can be found in Fig. S15 and S16.

GC-MS chromatograms indicated that the PA-H_2_O_2_ treatment decomposed cured epoxy resins into compounds with retention times of 7-9 and 12-14 minutes, as evidenced by distinct peaks A and B in BSA and control samples (Fig. 6a and S17). Notably, peak B was absent in samples treated with ItL-03, while peak A was additionally absent when combined with ABTS (Fig. S18). In ItL-03/ABTS samples, peak C appeared, and an additional peak D was detected after 24 hours (Fig. 6a). Peaks A and B, attributed to *m/z* 178 and 166, respectively, may have been degraded by laccase and could originate from NNBT or fractions of RTM6 (Fig. 6b). The ion at *m/z* 178 could possibly have been deprotonated to form the ion at *m/z* 177, corresponding to Peak C, as they exhibited similar ionization patterns. Alternatively, these ions may represent distinct molecules with nearly identical masses, such as the *m/z* pairs of 166-165 proposed for peaks B and D. Additional species at *m/z* 178, 177, 166, and 165 are proposed in Fig. S19 and S20. Furthermore, the ItL-03/ABTS treatment produced a prominent peak E, corresponding to *m/z* 144 (Fig. S21). This peak is likely a fragment of degraded ABTS, as it exhibited ionization patterns consistent with its mass fraction.

**Figure 6.**
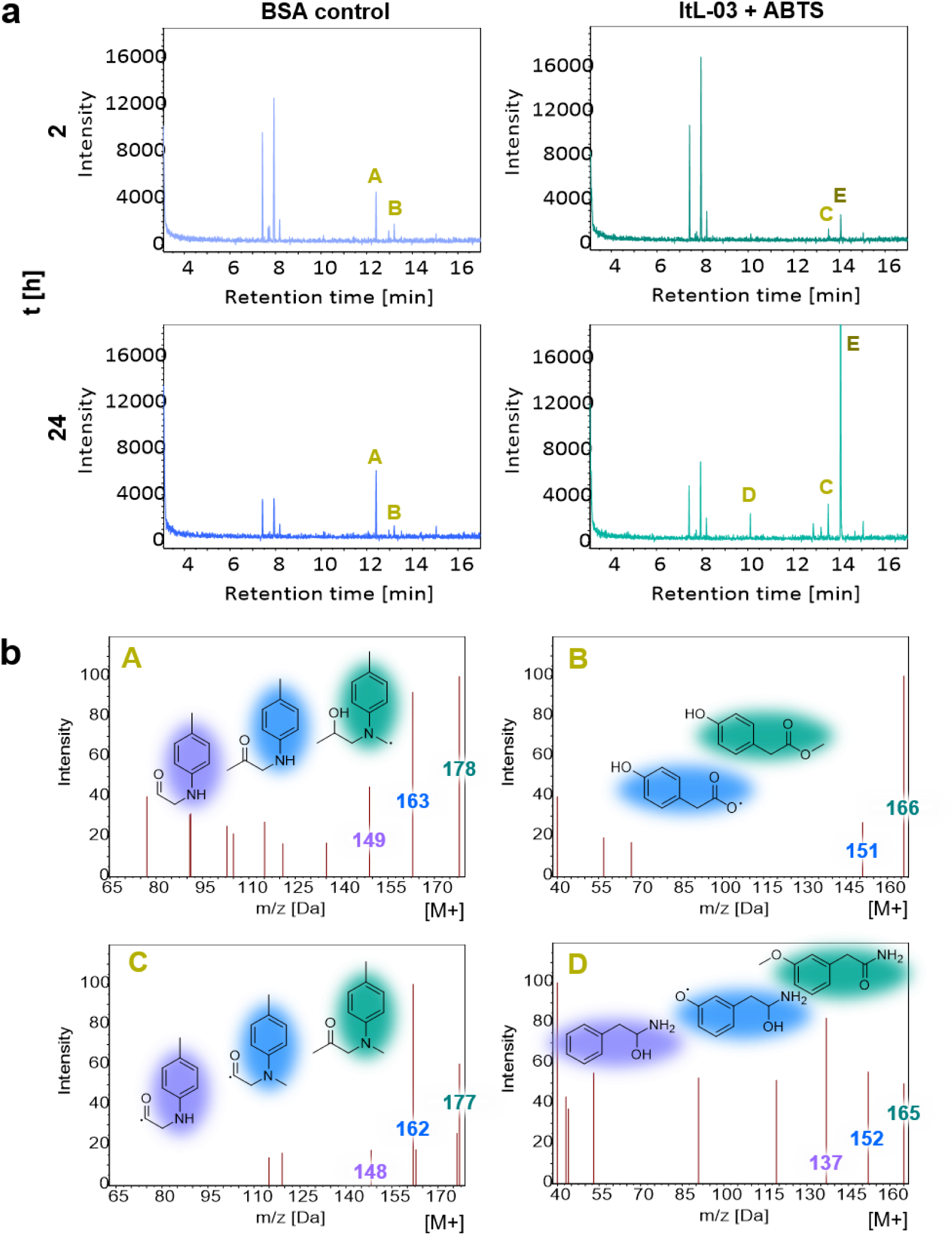
GC-MS analysis of extracted epoxy from chemo-enzymatic degradation using PA-H_2_O_2_ and laccase ItL-03 on RTM6. (a) GC-MS chromatogram presenting the degradation activity of ItL-03 with ABTS, compared to BSA control after 2 and 24 hours. The peaks of interest are indicated with A-E. (b) Proposed species derived from RTM6 epoxy that may be degraded by ItL-03, based on mass spectra of peaks A and B, with additional peaks C and D emerging in the ItL-03/ABTS samples after 2 and 24 hours. Chemical structures are color-highlighted to indicate different fragment ions *m/z*. Supporting data can be found in Fig. S17 and S18 and additional species related to peaks A-E are presented in Fig. S19-21.

Since the PA-H_2_O_2_ treatment of RTM6 from the same batch did not completely decompose the amine epoxy, any remaining epoxy powder was also tested for laccase ItL-03 activity. Before enzyme treatment, FTIR spectra showed distinct band patterns between non-treated (RTM6) and pre-treated (PT-RTM6) epoxy powders, indicating that PA-H_2_O_2_ treatment induced changes in the functional groups of epoxy resins (Fig. S22). The C=O stretching band (around 1,750 cm^-1^) was present only in PT-RTM6 and absent in RTM6, further confirming the oxidative effect of the pre-treatment on the DGEBF-based composites (Fig. 4c).

Subsequently, both RTM6 and PT-RTM6 were subjected to enzymatic treatment with ItL-03 in the presence of different mediators: ABTS, syringol, and TEMPO. For RTM6 samples, there was a significant change in the peak intensity of the C–N stretching associated with aromatic amines (1,210-1,180 cm^-1^) in the presence of enzyme (Fig. 7a). However, this effect resembled that of the BSA control, suggesting that the band shifts may be due to water incubation rather than enzymatic activity (Fig. S23). Epoxy resin is prone to absorb water and swell, potentially leading to structural degradation (Walter et al., 2013). This is not the case for the ItL-03/DMP, which exhibited slight changes that differed from the control. For PT-RTM6, epoxy functional groups, such as C–N stretching in aromatic (around 1,200 cm^-1^) and primary (1,085-1,050 cm^-1^) amines, were less pronounced in the samples treated with ItL-03 compared to those without (PT-RTM6) (Fig. 7b). A more significant effect was observed in the presence of mediators (Fig. S24). This suggests that pre-treatment facilitates enzyme access to the embedded epoxy functional groups in the polymer, particularly the amine functionalities, allowing for further oxidation.

**Figure 7.**
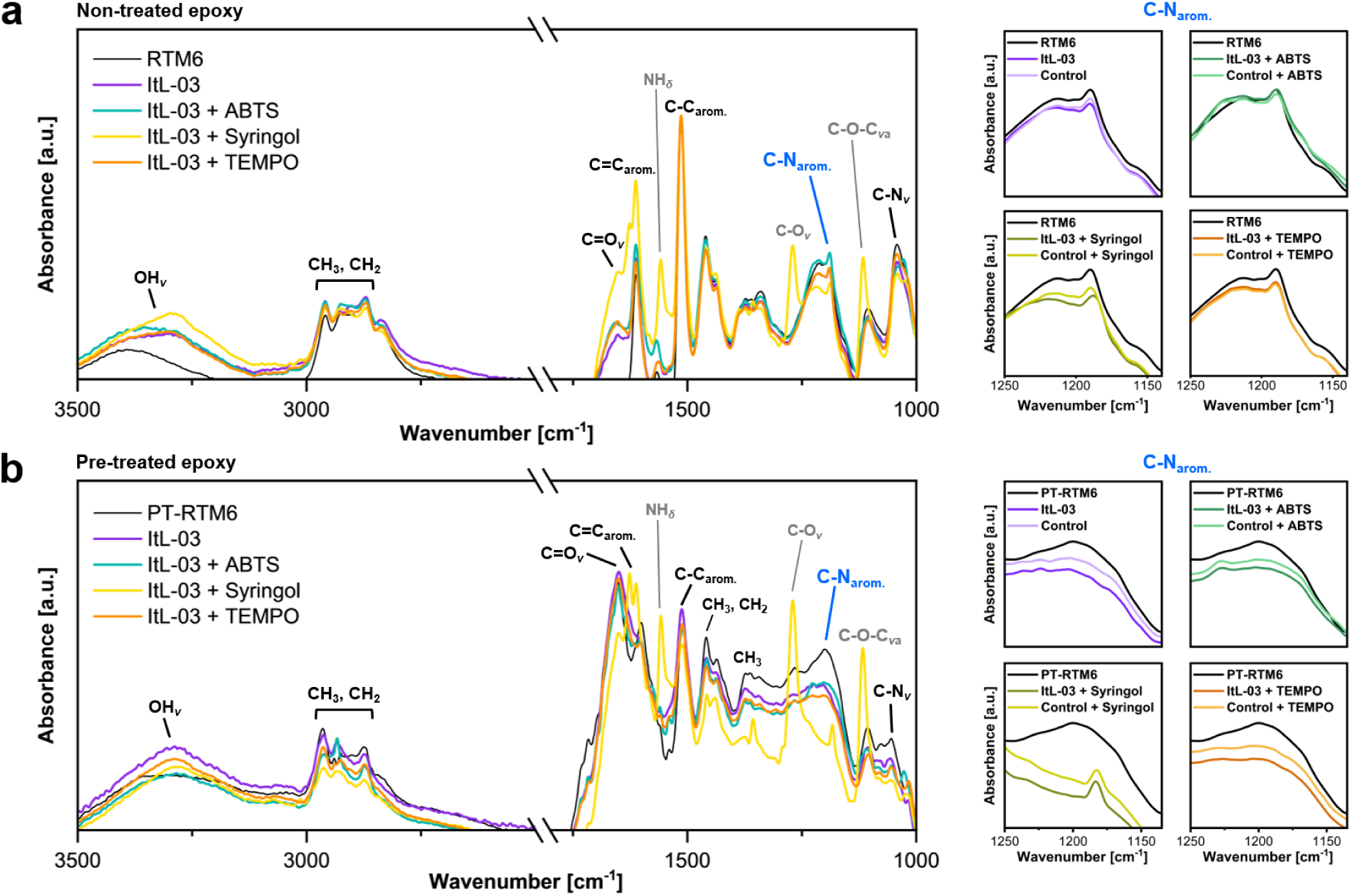
FTIR analysis of (a) RTM6 epoxy powder and (b) PA-H_2_O_2_ treated residual epoxy powder (PT-RTM6), focusing on the C-N_arom_. peaks highlighted in blue and shown on the right. Both RTM6 and PT-RTM6 were subsequently treated for 5 days with laccase ItL-03 and different mediators: ABTS, syringol, and TEMPO. The samples labeled ‘RTM6’ and ‘PT-RTM6’ were not subjected to enzyme treatment and the control refers to the BSA incubation of the epoxy instead of laccase. The corresponding functional groups are indicated. *v* — stretching; *v*a — asymmetric stretching; *δ* — deformation. Other functional group peaks, in comparison to BSA, are shown in Fig. S23 and S24.

## 4. Discussion

In recent years, the search for sustainable waste management solutions has increasingly focused on biodegradation of recalcitrant synthetic polymers like epoxy resins. To date, no microorganism or enzyme has been clearly shown to effectively degrade cured epoxy polymers (PAZy database; Buchholz et al. (2022)). Our study represents an early step forward by demonstrating that a two-step process involving PA-H_2_O_2_ pre-treatment followed by laccase ItL-03 can modify RTM6 epoxy resins and release defined monomers over time.

The studies by Dolz et al. (2022) represent one of the first investigations into NNBT, demonstrating that it can be degraded by unspecific peroxygenases (UPOs) through N-dealkylation. Our study shows that bacterial laccases can also degrade NNBT and expand the scope to include other tertiary amine epoxy substrates, BBCM and BMAP, derived from Hexflow® RTM6, through an N-dealkylation activity.

Laccases readily degrade C–N bonds, but have minimal effect on modifying the surface of epoxy polymer, without pre-treatment. The steric hindrance from the densely packed, cross-linked structure likely limits enzyme access to certain regions within the molecule (Escayola et al., 2024; Pinter et al., 2012). For recalcitrant polymers like epoxy, pre-treatment with peracid (or PA-H_2_O_2_) generates acyloxy and hydroxyl radicals that oxidize C–O bonds into carbonyls and cleave C–N bonds, breaking down the epoxy cross-linking (Das et al., 2018). Thus, laccase more evidently target amine functionalities, reducing C–N, CH_3_, CH_2_, and C–O–C linkages.

Modifications of these epoxy functional groups have been reported in marine bacteria, which decrease the corrosion resistance of epoxy-coated steel after one month in seawater. *Pseudomonas aeruginosa* reduced C–O–C and C–O groups (S. Zhang et al., 2023), while *Bacillus flexus* broke down aromatic rings and epoxy groups (Deng et al., 2019). A soil bacterial consortium used epoxy as the main carbon source, reducing C–N, CH_3_, CH_2_, and C–O groups, with 34% polymer weight loss (Pardi-Comensoli et al., 2022).

Colonization and environmental factors, such as seawater, can accelerate epoxy oxidation and hydrolysis (Da Costa et al., 2018; Dang & Lovell Charles, 2015), mimicking pre-treatment that prepares substrates for enzyme activity. Bacteria and fungi colonizing epoxy polymers (Gu et al., 1997; Wang et al., 2016) often possess oxidoreductases (EC1), such as laccases, peroxidases, monooxygenases, and alcohol dehydrogenases (Kumar & Chandra, 2020; Naveed et al., 2025; Padayachee et al., 2020), that have potential to degrade non-hydrolyzable plastics like PE, PP, PS, and PVC (Chow et al., 2023; Mohanan et al., 2020).

Other studies have shown that combining pre-treatment with enzymes enhances degradation of non-hydrolyzable polymers. Branson et al. (2023) demonstrated that glycolyzing polyether polyurethane foams at 200 °C with diethylene glycol (DEG) and 1% tin (II)-2-ethylhexanoate produced low-molecular-weight (LMW) dicarbamates, which were hydrolyzed into aromatic diamines with urethanase UMG-SP-2, achieving a 65% conversion in 24 hours. Similarly, pre-treating LMWPE with m-chloroperoxybenzoic acid (mCPBA) and ultrasonication improved enzyme accessibility, resulting in approximately 27% polymer conversion and releasing medium-chain products, such as aliphatic carboxylic acids (Oiffer et al., 2024).

Combining PA-H_2_O_2_ treatment with laccase ItL-03 has shown promise in enhancing the degradation of epoxy polymers compared to using enzymes alone. However, this approach often results primarily in surface modification rather than significant degradation throughout the material. While this approach improves enzyme accessibility, the degradation level may not meet industrial requirements. This suggests an area for further development.

Expanding enzyme screening to include various epoxy types, such as secondary amines (R–NH– CH2–), carboxyl esters (R–(C=O)–O–R’), ethers (R–O–R’), and thiol (R–S–R’) linkages (Rashid et al., 2024), and larger model substrates would improve our understanding of enzymatic mechanisms, particularly for bacterial laccases with high redox mediators. Though propionic acid can be biologically produced through microbial fermentation, such as with *Propionibacterium freudenreichii* and *P. acidipropionici* (Gonzalez-Garcia et al., 2017; Ranaei et al., 2020), and recovered for reuse through methods like microchannel distillation (Lu et al., 2023; Singh et al., 2021), minimizing pre-treatment is important to ensure adequate enzyme access while adhering to greener practices.

Effective recycling of epoxy resins necessitates a multifaceted approach that incorporates physical, chemical, and enzymatic methods. While significant challenges remain, our study offers an innovative biological concept for epoxy waste management. Continued research will be essential to transform this potential into practical, scalable recycling methods.

## DATA AVAILABILITY

Metagenomic sequencing data were deposited in the Integrated Microbial Genomes (IMG) database under project accession number Gp0109785. All data generated or analyzed during this study are included in this manuscript and its supplementary information files, including Supplementary Data. The corresponding author can provide further data that support the findings of this study upon reasonable request.

## AUTHOR CONTRIBUTIONS

S.W., P.P.G., and W.R.S designed the study and contributed to manuscript writing. A.S. performed enrichments and isolation of metagenomic DNA. S.W. and A.S. performed cloning. S.W. conducted experiments and analyzed data. L.K. synthesized the amine epoxy powder. L.K., A.M.R, and A.L. contributed to FTIR analysis of CFRPs. C.C. provided EDX results. W.R.S received funding. All authors revised and accepted the manuscript.

## ACKNOWLEDGEMENTS

We would like to thank Phillip Schlottau from the Department of Physics, for the assistance with laser cutting on epoxy. We acknowledge Elke Wölken from the Microscopy Department, with support from SEM, as well as the Core Facility Mass Spectrometry, part of the Technology Platform Mass Spectrometry, particularly Gaby Graack and Erik Mordhorst, for their support with mass spectrometric analysis. We extend our thanks to Siraphat Weerathaworn from the Institute of Physical Chemistry for FTIR assistance. All contributors are from the University of Hamburg. The epoxy substrate BMAP was kindly provided by Prof. Nigel Scrutton from the Manchester Institute of Biotechnology at The University of Manchester (UK). We are grateful to Max Kolb from Airbus Defense & Space GmbH, X-Labs (Munich, Germany) for performing SEM/EDX analysis on the pre-treated CFRPs.

