## Supporting Information for "A Combined Chemo-Enzymatic Treatment for the Oxidation of Epoxy-Based Carbon Fiber-Reinforced Polymers (CFRPs)"

**Table S1.** Putative multicopper oxidase enzymes from *I. typographus* and other bacterial laccases known for their association with lignin and synthetic polymer degradation.

| Name | Accession number | Cluster | Phylogenetic affiliation | aa identity with CueO [%] | Copper binding motifs | | | | Reference |
|---|---|---|---|---|---|---|---|---|---|
| | | | | | *i* | *ii* | *iii* | *iiii* | |
| CueO | P36649* | γ-proteobacteria/C1 | *Escherichia coli* | 100.00 | **H**W**H** | **H**P**H** | **H**PF**H**I**H** | **HCH**LLE**H**EDTG**M** | 1 |
| ItL-01 | Ga0063521_10002204** | γ-proteobacteria/C1 | *Erwinia* sp. | 63.18 | **H**W**H** | **H**P**H** | **H**PF**H**I**H** | **HCH**LLE**H**EDTG**M** | This work |
| ItL-02 | Ga0063521_100014138** | γ-proteobacteria/C1 | *Dryocola clanedunensis* | 74.75 | **H**W**H** | **H**P**H** | **H**PF**H**I**H** | **HCH**LLE**H**EDTG**M** | This work |
| ItL-03 | Ga0063521_100024328** | γ-proteobacteria/C1 | *Rahnella* sp. | 63.96 | **H**W**H** | **H**P**H** | **H**PF**H**I**H** | **HCH**LLE**H**EDTG**M** | This work |
| ItL-04 | Ga0063521_1000001282** | γ-proteobacteria/C1 | *Providencia* sp. | 61.49 | **H**W**H** | **H**P**H** | **H**PF**H**V**H** | **HCH**LLE**H**EDTG**M** | This work |
| ItL-05 | Ga0063521_100021622** | γ-proteobacteria/C1 | *Morganella morganii* | 64.02 | **H**W**H** | **H**P**H** | **H**PF**H**I**H** | **HCH**LLE**H**EDTG**M** | This work |
| ItL-06 | Ga0063521_100028714** | γ-proteobacteria/C1 | *Rahnella* sp. | 63.51 | **H**W**H** | **H**P**H** | **H**PF**H**I**H** | **HCH**LLE**H**EDTG**M** | This work |
| hh01-218 | 2523500218** | β-proteobacteria/C2 | *Janthinobacterium* sp. | 38.37 | **H**W**H** | **H**P**H** | **H**PI**H**L**H** | **HCH**KSH**H**TMNA**M** | 2 |
| hh01-482 | 2523500482** | β-proteobacteria/C2 | *Janthinobacterium* sp. | 38.46 | **H**W**H** | **H**P**H** | **H**PI**H**I**H** | **HCH**KSH**H**TMNA**M** | 2 |
| sm3532 | 642693532** | γ-proteobacteria/C2 | *Stenotrophomonas maltophilia* | 23.91 | **H**W**H** | **H**S**H** | **H**PI**H**L**H** | **HCH**LLY**H**MEAG**M** | 3 |
| sm4696 | 642694696** | γ-proteobacteria/C2 | *Stenotrophomonas maltophilia* | 22.46 | **H**W**H** | **H**S**H** | **H**PI**H**L**H** | **HCH**LLY**H**MEAG**M** | 3 |
| ngr147 | 643825147** | α-proteobacteria/C2 | *Sinorhizobium fredii* | 23.04 | **H**W**H** | **H**P**H** | **H**PI**H**L**H** | **HCH**IIE**H**QKTG**M** | 4 |
| ngr578 | 643822578** | α-proteobacteria/C2 | *Sinorhizobium fredii* | 36.19 | **H**W**H** | **H**P**H** | **H**PI**H**M**H** | **HCH**KSH**H**TMNA**M** | 4 |
| ngr600 | 643822600** | α-proteobacteria/C2 | *Sinorhizobium fredii* | 33.67 | **H**W**H** | **H**S**H** | **H**PM**H**L**H** | **HCH**HLY**H**MNGG**M** | 4 |
| ngr688 | 643822688** | α-proteobacteria/C2 | *Sinorhizobium fredii* | 36.19 | **H**W**H** | **H**P**H** | **H**PI**H**M**H** | **HCH**KSH**H**TMNA**M** | 4 |
| LMCO2 | AII11185.1 | Actinomycetes/C2 | *Rhodococcus opacus* | 26.38 | **H**W**H** | **H**P**H** | **H**PM**H**L**H** | **HCH**NLY**H**GEAG**M** | 5 |
| LMCO3 | AII11221.1 | Actinomycetes/C2 | *Rhodococcus opacus* | 31.19 | **H**L**H** | **H**A**H** | **H**TM**H**F**H** | **HCH**VGPLAE**H** | 5 |
| LfLAC3 | WP_193831439.1 | Bacilli/C1 | *Lysinibacillus fusiformis* | 25.48 | **H**L**H** | **H**D**H** | **H**PI**H**L**H** | **HCH**FLE**H**EDHD**M** | 6 |
| RrLAC2 | WP_318283942.1 | Actinomycetes/C2 | *Rhodococcus ruber* | 25.69 | **H**F**H** | **H**S**H** | **H**PM**H**V**H** | **HCH**NAY**H**QEAG**R** | 6 |

*UniProt, **IMG, C: clade, *i*-*iiii* indicate amino acids involved in copper binding motifs.

**Table S2.** Bacterial strains, plasmids, and primers used in this work.

| Strain | Characteristics | Source/Reference | |
|---|---|---|---|
| *E. coli* DH5α | *sup*E44, Δ*lac*U169 (Φ80 *lac*ZΔM15) *hsd*R17 *rec*A1 *end*A1 *gyr*A96 *thi*-1 *rel*A1 | Invitrogen, Germany | |
| *E. coli* BL 21 (DE3) | F- *omp*T *hsd*SB (rB- mB-) *gal dcm*, (DE3) | Novagen/Merck, Germany | |
| *E. coli* T7 SHuffle | F´ *lac*, *pro*, *lacI*$^{q}$ / Δ(*ara-leu*)7697 *ara*D139 *fhu*A2 *lac*Z::*T7 gene1 Δ(phoA)PvuII phoR ahpC* galE (or U) galK λatt*::pNEB3-r1-cDsbC (Spec$^{R}$, *lacI*$^{q}$) ΔtrxB rpsL150(Str$^{R}$) Δgor Δ(malF)3 | New England BioLabs, Germany | |
| **Vector** | **Characteristics** | **Source/Reference** | |
| pET21a::CueO | MCO CueO from *E. coli* in pET21a with His-tag | This study | |
| pET21a::ItL-01 | Potential MCO ItL-01 from *I. typographus* metagenome in pET21a with His-tag | This study | |
| pET21a::ItL-02 | Potential MCO ItL-02 from *I. typographus* metagenome in pET21a with His-tag | This study | |
| pET21a::ItL-03 | Potential MCO ItL-03 from *I. typographus* metagenome in pET21a with His-tag | This study | |
| **Primer** | **Sequence 5' → 3'** | **GC Content [%]** | **Length [nt]** |
| pET_for | ATATAGGCGCCAGCAACC | 56 | 18 |
| MCO_HindIII_rev | CGTCAGAAGCTTGCTCCATGCAGAGAGCTC | 57 | 30 |
| T7_prom | TAATACGACTCACTATAGGG | 40 | 20 |
| T7_term | CTAGTTATTGCTCAGCGGT | 47 | 19 |

**Table S3.** Kinetic constants of bacterial laccases for ABTS substrate.

| Enzymes | $K_{m}$ [mM] | $K_{cat}$ [min$^{-1}$] | $K_{cat}/K_{m}$ [min$^{-1}$ mM$^{-1}$] |
|---|---|---|---|
| CueO | 2.39 ± 0.23 | 0.27 ± 0.03 | 0.11 ± 0.01 |
| ItL-01 | 4.50 ± 1.42 | 0.10 ± 0.01 | 0.02 ± 0.01 |
| ItL-02 | 3.64 ± 0.26 | 0.80 ± 0.01 | 0.22 ± 0.01 |
| ItL-03 | 7.76 ± 0.88 | 3.04 ± 0.24 | 0.39 ± 0.02 |

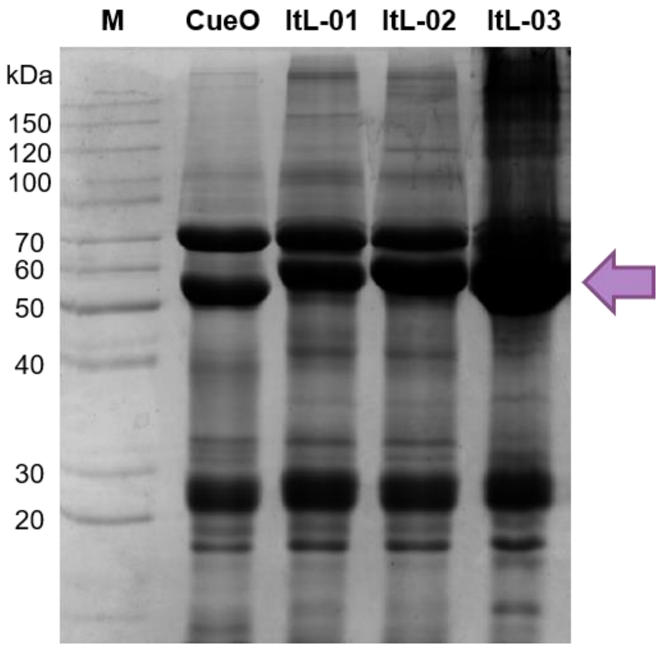


**Figure S1.** SDS-PAGE analysis shows that CueO and ItL-01-03 were successfully expressed and purified using Ni-NTA chromatography. M indicates an unstained protein ladder (ThermoScientific, Bremen, Germany), with the arrow pointing to the expected bands of the laccases, around 55-65 kDa. Additional non-specific bands observed at 25 kDa and 70 kDa reflect the complexity of the protein sample, as the *E. coli* host expressed these bands concurrently.

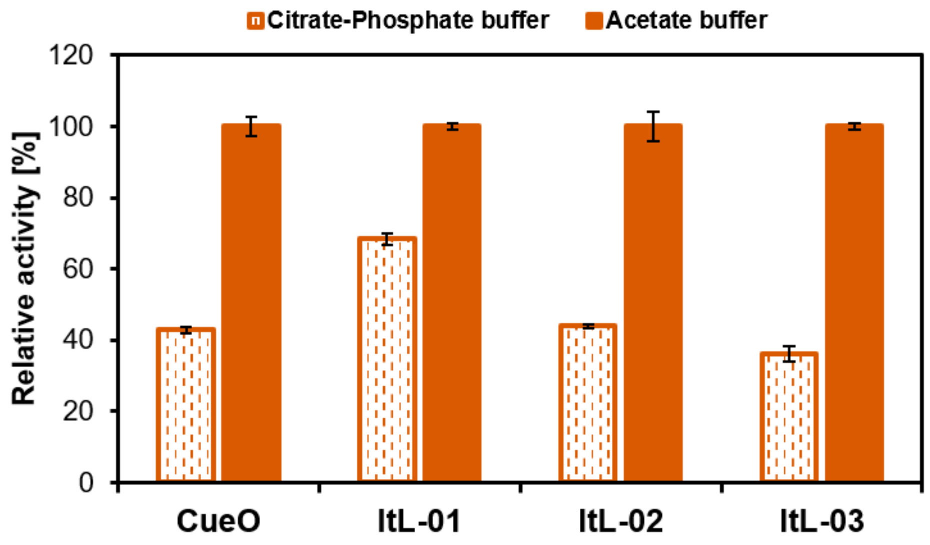


**Figure S2.** Buffer preference: laccases prefer acetate buffer, showing a 30-50% increase. Error bars indicate the standard deviation (*n* = 3).

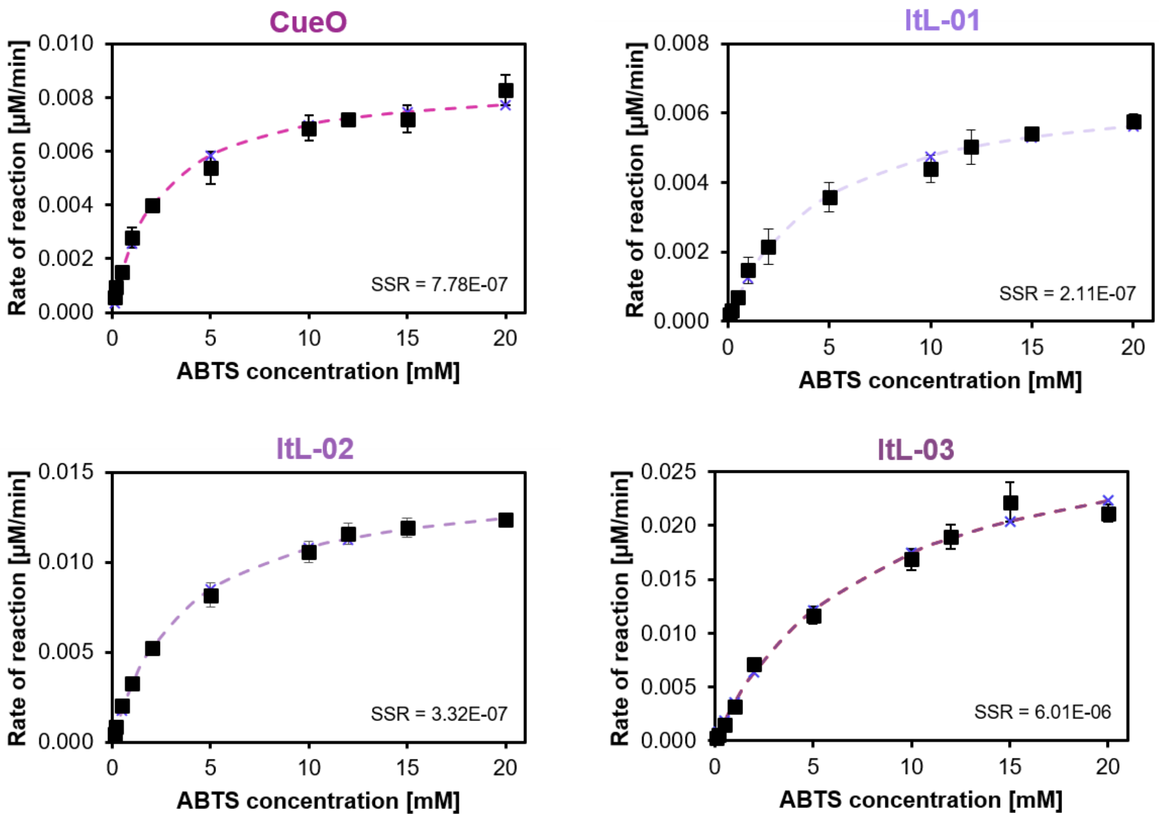


**Figure S3.** Kinetic constants of laccases for ABTS oxidation. Kinetic parameters were estimated by fitting the data to the Michaelis-Menten equation using Solver, with the corresponding sum of squared residuals (SSR). Error bars indicate the standard deviation ($n$ = 3).

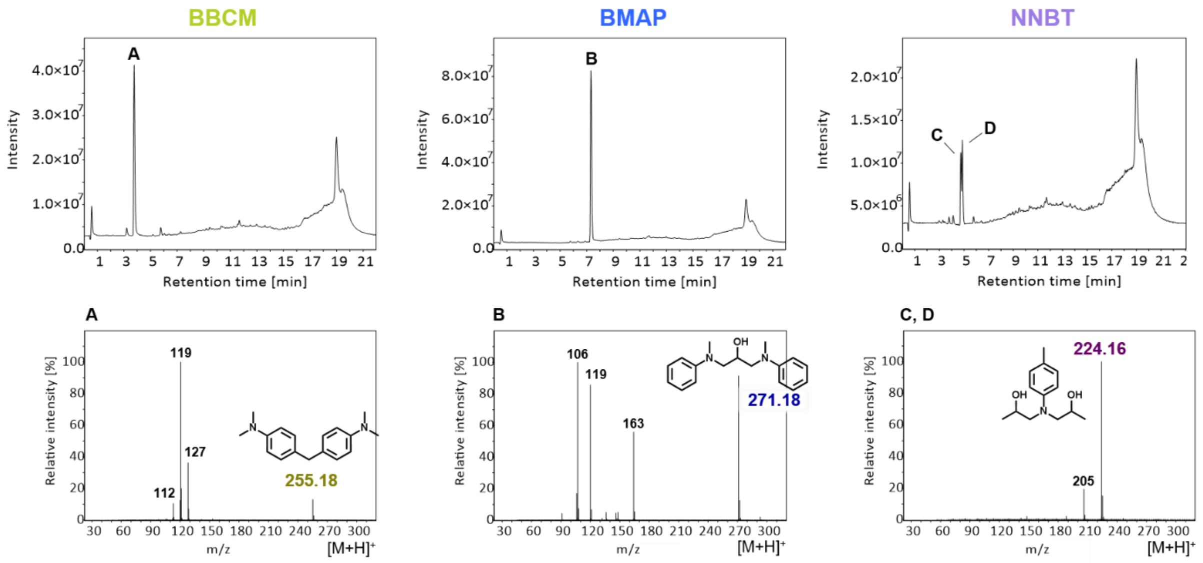


**Figure S4.** LC-MS analysis identifying epoxy model compounds in positive mode, based on their retention times, with specific signals corresponding to their exact masses: BBCM (A): 3.8 min, m/z 254, BMAP (B): 7.3 min, m/z 270, NNBT (C, D): 4.7 and 4.8 min, m/z 223.

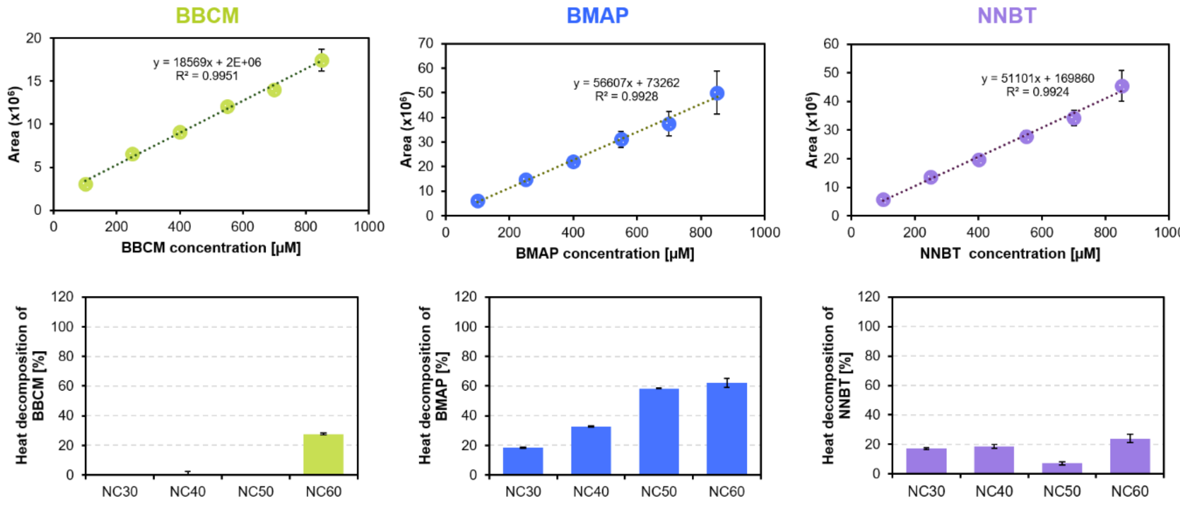


**Figure S5.** (top) Calibration curve to determine concentrations of epoxy model substrates using LC-MS. (bottom) The effect of temperature (30-60 °C) on the decomposition of epoxy models (BBCM, BMAP, NNBT) was evaluated after 2 hours as a negative control in the absence of enzymes. Error bars indicate the standard deviation (*n* = 3).

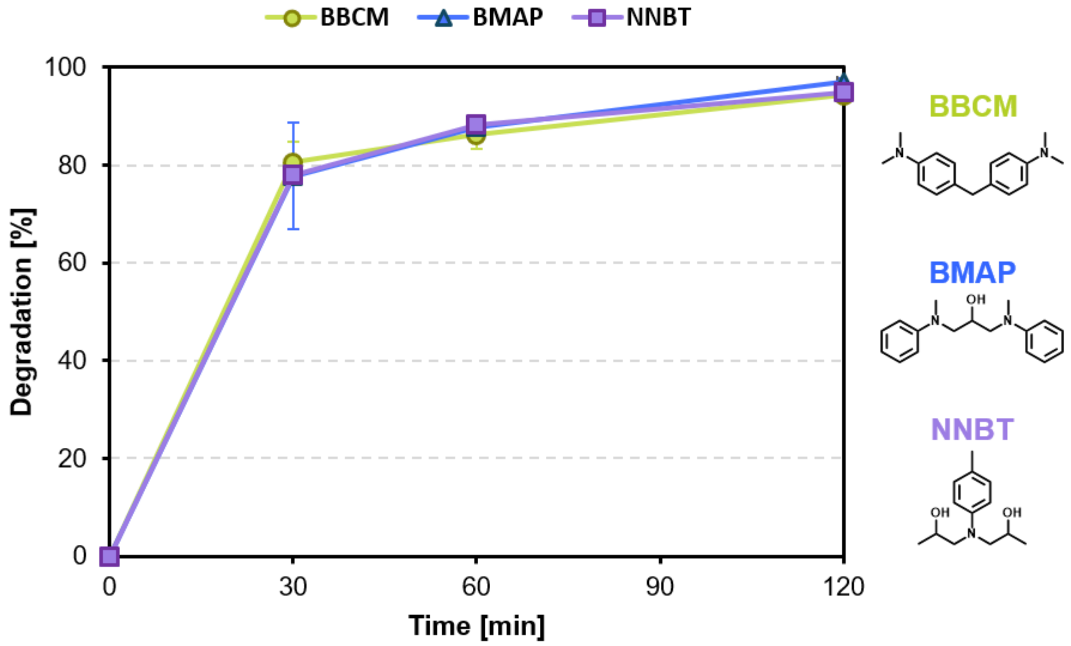


**Figure S6.** Degradation activity of ItL-03 on epoxy scaffolds of amine epoxy: BBCM, BMAP, and NNBT. Measurements using LC-MS were taken after 0, 30, 60, and 120 minute(s) of incubation at 50 °C in 0.1 M acetate buffer pH 4.0. The initial observation ($t_0$) defines the starting amount of epoxy, set at 100%. The degradation rate is calculated by subtracting the percentage of remaining epoxy from 100%. ItL-03 reaches approximately 80% degradation of these substrates within 30 minutes and nearly complete degradation after a duration of 2 hours. Error bars indicate the standard deviation (*n* = 3).

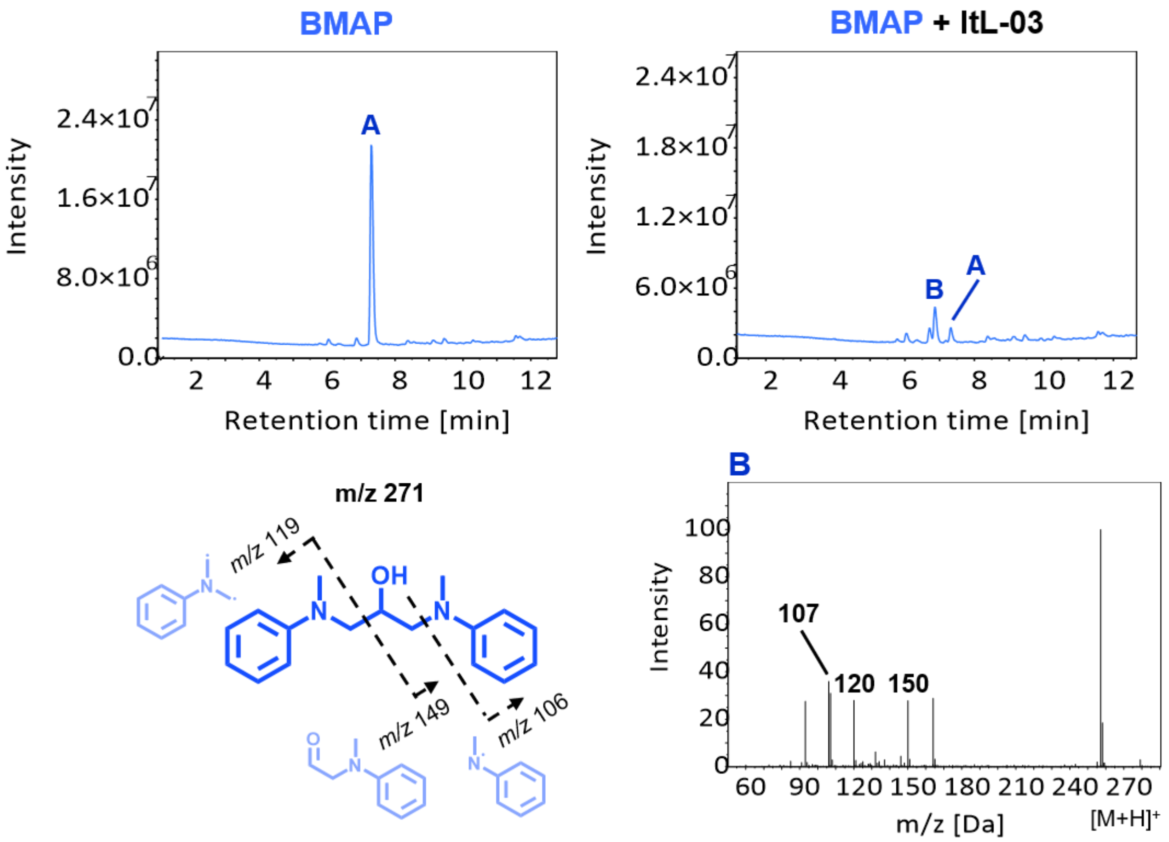


**Figure S7.** LC-MS chromatogram of the reaction involving laccase ItL-03 and BMAP. (A) The mass of BMAP is observed at 7.3 minutes. (B) The mass spectrum of the N-dealkylation product reveals a peak that appears at 6.7 minutes.

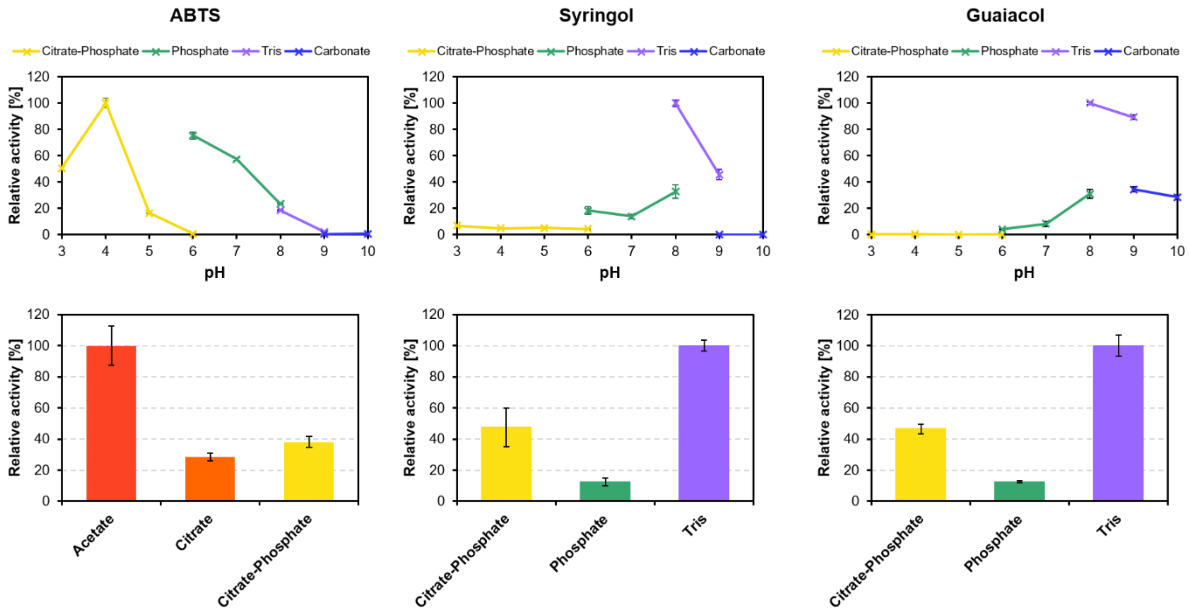


**Figure S8.** (top) pH profile of laccase ItL-03 toward different mediators: ABTS, syringol, and guaiacol. The enzyme exhibits maximum activity at pH 4.0 for ABTS, while displaying a preference for pH 8.0 for both syringol and guaiacol. (bottom) Buffer preference of ItL-03 with each mediator. For ABTS, ItL-03 shows the highest activity in acetate buffer, while for syringol and guaiacol, optimal activity is observed in tris buffer. Error bars represent the standard deviation ($n$ = 3).

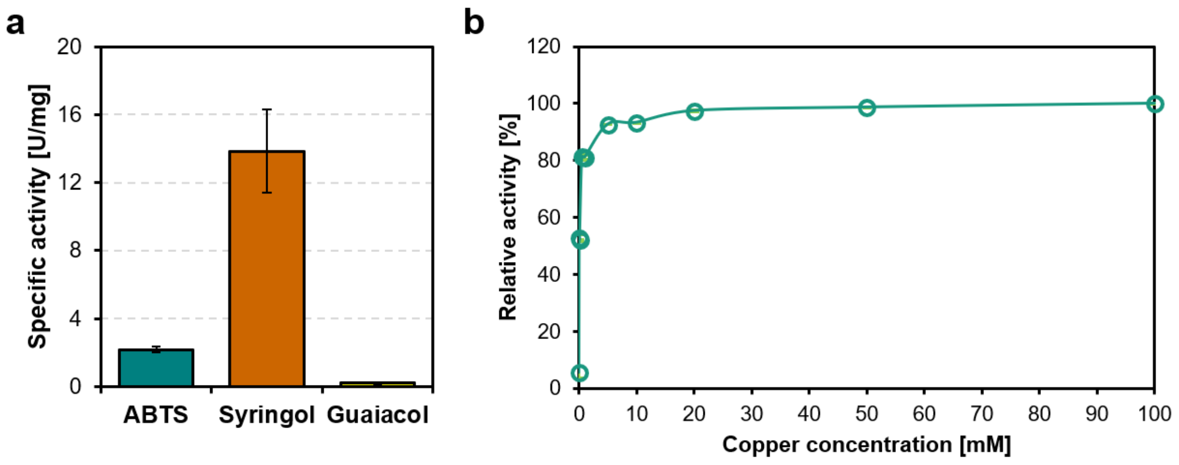


**Figure S9.** (a) Specific activity of laccase ItL-03 toward ABTS, syringol, and guaiacol. Each substrate was evaluated in its respective optimal buffer and pH environment (Fig. S8). Laccase activity revealed that syringol had the highest activity, about four times greater than ABTS, while guaiacol showed relatively low activity. (b) Optimal copper ions concentration for ItL-03. Enzyme activity significantly increases with elevated $Cu^{2+}$, saturating at around 5 mM. Error bars represent the standard deviation ($n$ = 3).

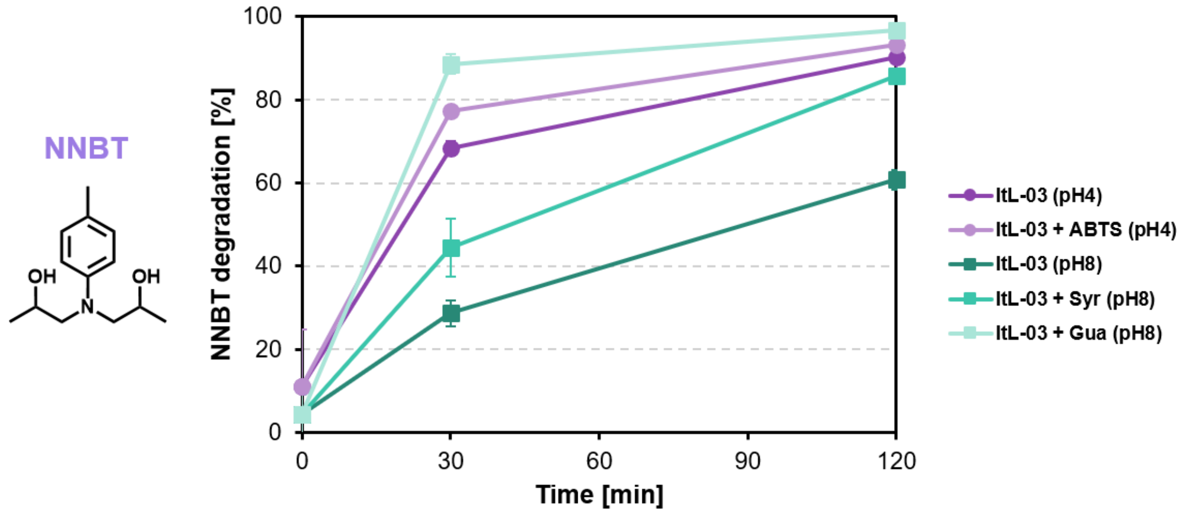


**Figure S10.** Degradation activity of ItL-03 on NNBT (epoxy model) in the presence and absence of different mediators: ABTS, syringol (Syr), and guaiacol (Gua). Measurements using LC-MS were taken after 0, 30, and 120 minute(s) of incubation. The initial observation ($t_0$) defines the starting amount of epoxy, set at 100%. The degradation rate is calculated by subtracting the percentage of remaining epoxy from 100%. ItL-03/guaiacol exhibits higher activity than ItL-03/ABTS at 30 minutes, although both achieve similar levels of activity, over 90%, after 2 hours. Error bars indicate the standard deviation ($n$ = 3).

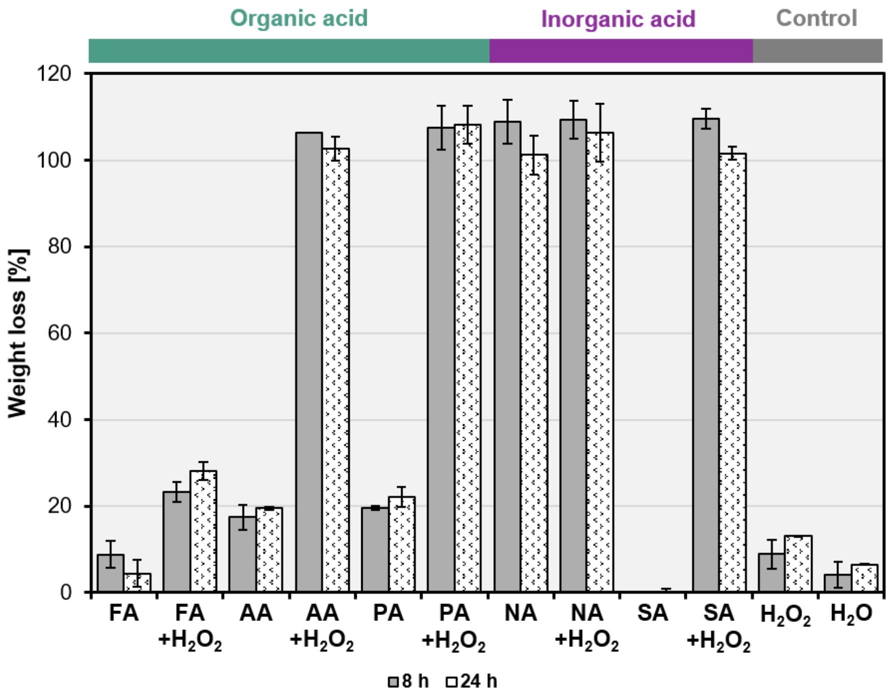


**Figure S11.** Weight loss of the resin mass in eCFRPs treated with organic acids, with and without hydrogen peroxide ($H_2O_2$), was compared to inorganic acids and controls ($H_2O$ and $H_2O_2$). The acids tested included formic acid (FA), acetic acid (AA), propionic acid (PA), nitric acid (NA), and sulfuric acid (SA). Composites were incubated in a 95:5 acid-to-peroxide ratio at a 9 M, 65 °C, and 200 rpm for 8 and 24 hours. Error bars indicate the standard deviation (*n* = 3). The effectiveness of these acid-peroxide treatments resulted in complete fiber recovery, probably causing a small loss of carbon fibers during washing, which could explain the weight loss values exceeding 100%.

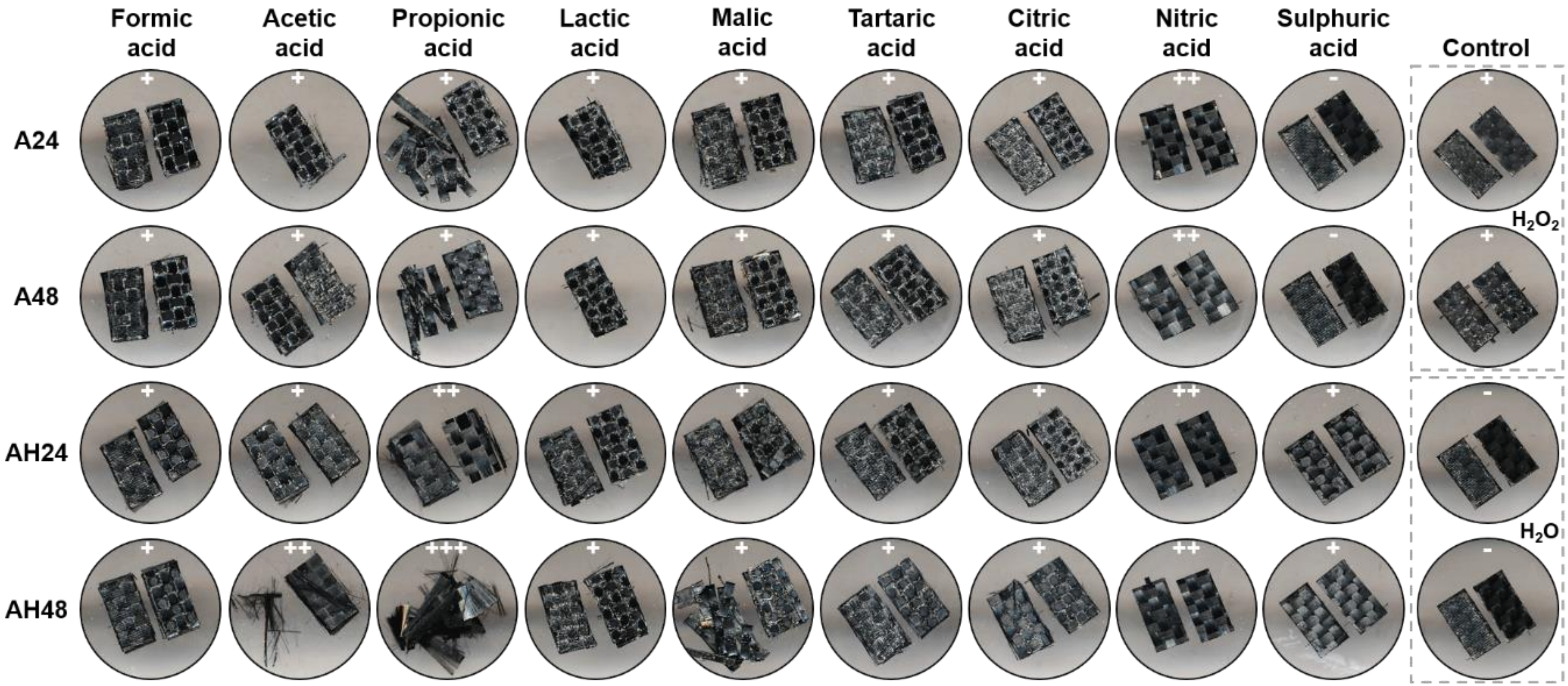


**Figure S12.** Morphological changes in eCFRPs after 24 and 48 hours of pre-treatment, with only acid (A) and with acid and $H_2O_2$ (AH), compared to controls ($H_2O$ and $H_2O_2$). The composites were incubated with a 95:5 acid-to-peroxide ratio at a 5 M, 65 °C and 200 rpm. +, ++, and +++ represent the degree of carbon fiber exposure, ranging from low to high. – indicates no change observed.

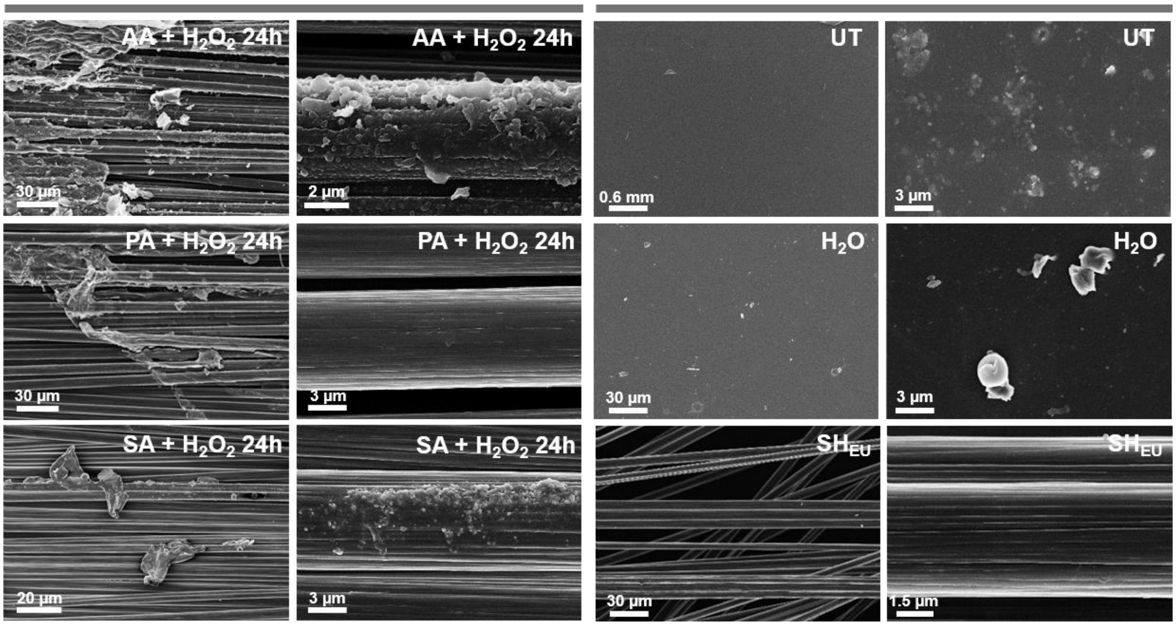


**Figure S13.** SEM images of eCFRPs after 24-hour treatments with acid (A) and hydrogen peroxide ($H_2O_2$). These results are compared to those from European protocol ($SH_{EU}$, DIN EN 2564:2018) and controls, including an untreated sample (UT) and $H_2O$. Scale bars indicate size in micrometers (µm) and millimeters (mm).

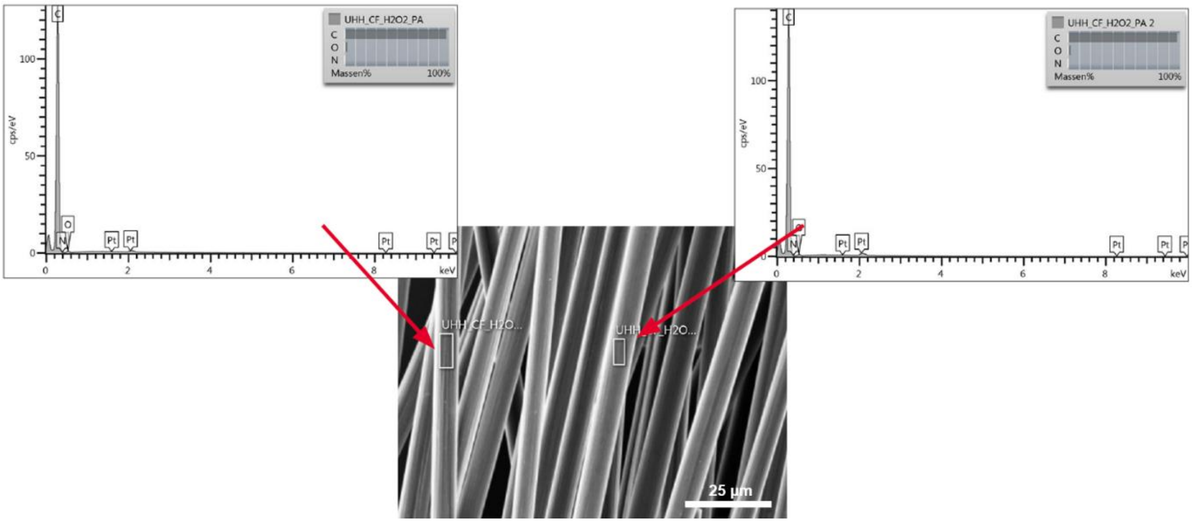


**Figure S14.** EDX spectra of carbon fibers (CFs) recovered after propionic and $H_2O_2$ pre-treatment. Two locations on the CFs (indicated by red arrows) were examined for elemental composition via EDX. The predominant element detected is carbon (C), with a very small percentage of oxygen (O), and no nitrogen (N) atom was detected. Scale bars indicate size in micrometers (µm).

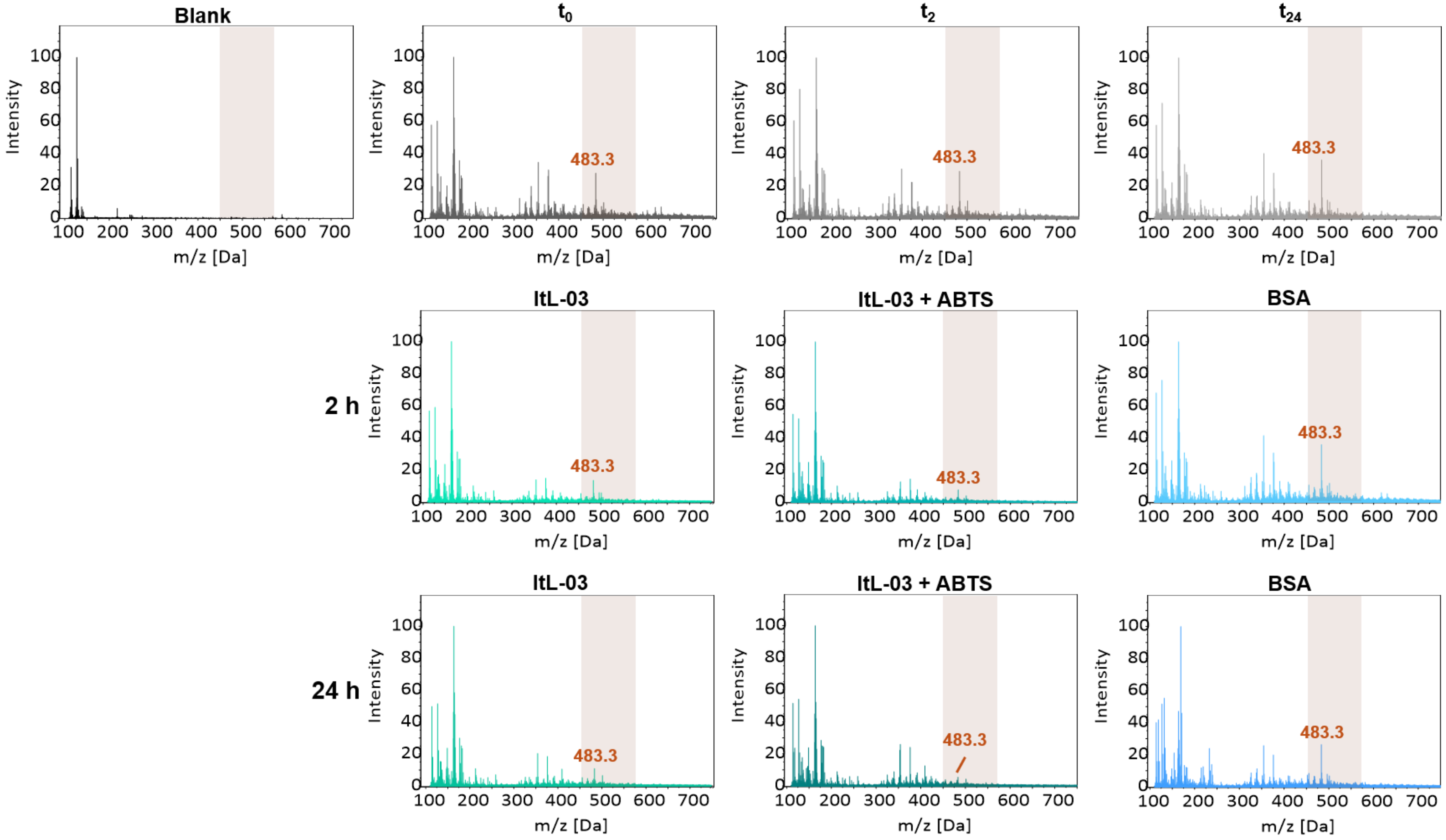


**Figure S15.** ESI-MS spectra in positive mode of the degradation activity of ItL-03 with or without ABTS, compared to BSA on extracted epoxy after pre-treatment (PA-$H_2O_2$). Measurements were collected after 0, 2, and 24 hours. $t_0$, $t_2$, and $t_{24}$ represent controls without enzyme at the beginning, between, and end of the incubation period. The ions *m/z* 483.3 was mostly absent in the laccase sample but remained present in the control.

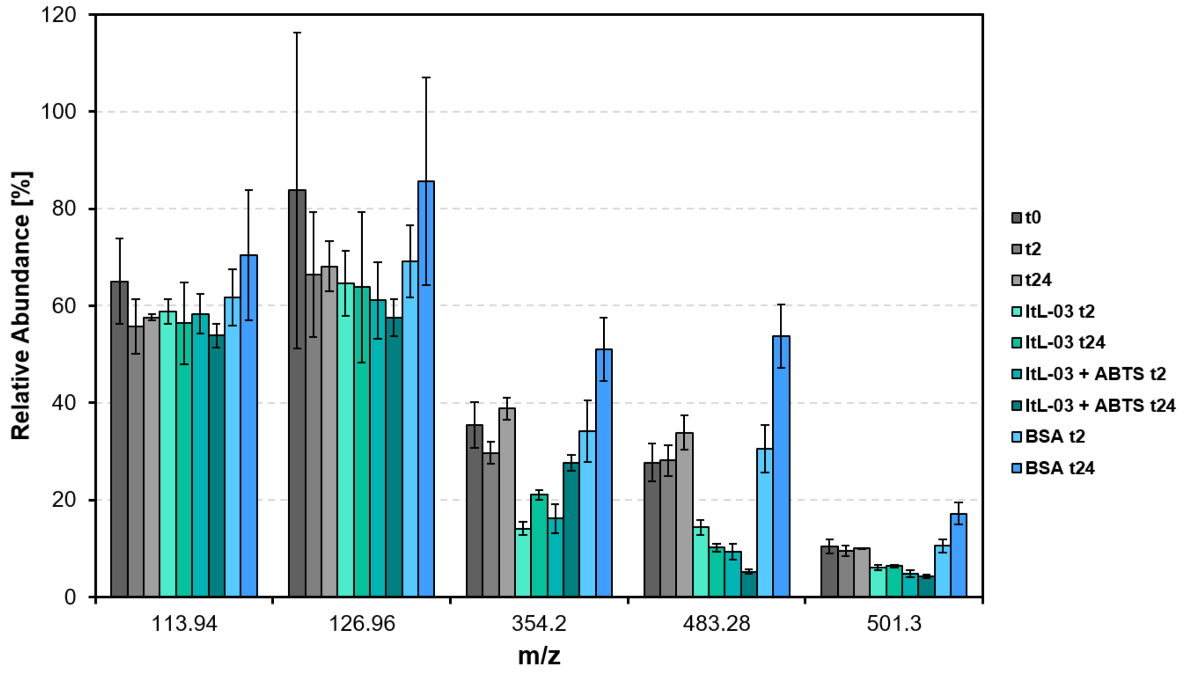


**Figure S16.** The relative abundance of *m/z* values of interest from the same experiment in 'Fig. S15' is shown. The spectra were normalized using the *m/z* value at 163.9 as a reference peak. $t_0$, $t_2$, and $t_{24}$ represent controls without enzyme at the beginning, between, and end of the incubation period. The abundance of the five ions is lower in the samples treated with the ItL-03 compared to the controls, particularly at *m/z* 354.2 and 483.3. Error bars represent the standard deviation ($n$ = 3).

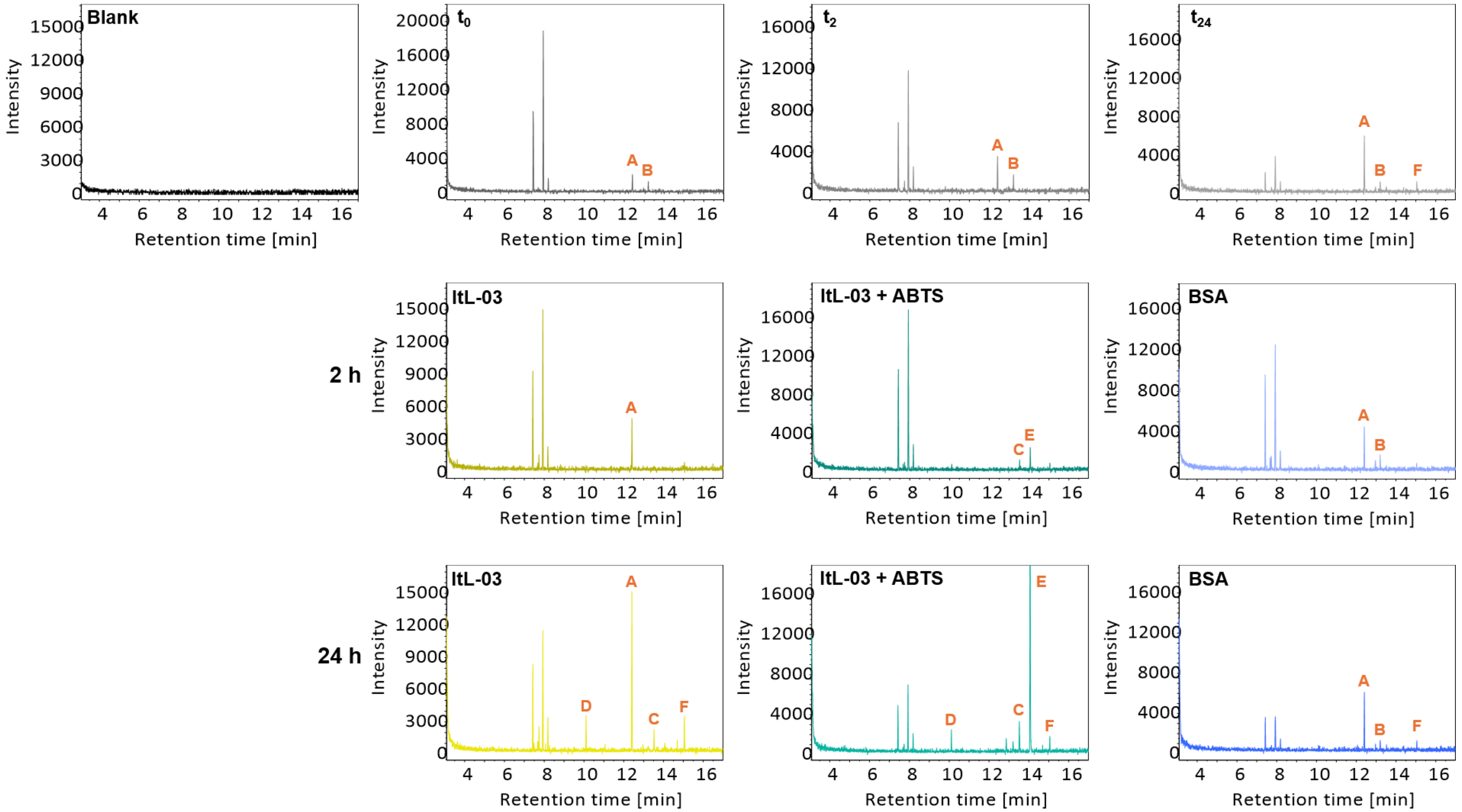


**Figure S17.** GC-MS chromatogram of the degradation activity of ItL-03 with or without ABTS, compared to BSA on extracted epoxy after pre-treatment (PA-$H_2O_2$). Measurements were collected after 0, 2, and 24 hours. $t_0$, $t_2$, and $t_{24}$ represent controls without enzyme at the beginning, between, and end of the incubation period. The peaks of interest are indicated with A-F. Peaks A and B were observed in BSA and control samples but were absent in the ItL-03/ABTS samples. Peaks C and D were detected in ItL-03 samples after 24 hours, while peak E was only present in samples containing ABTS. Peak F was detected in all samples after 24 hours.

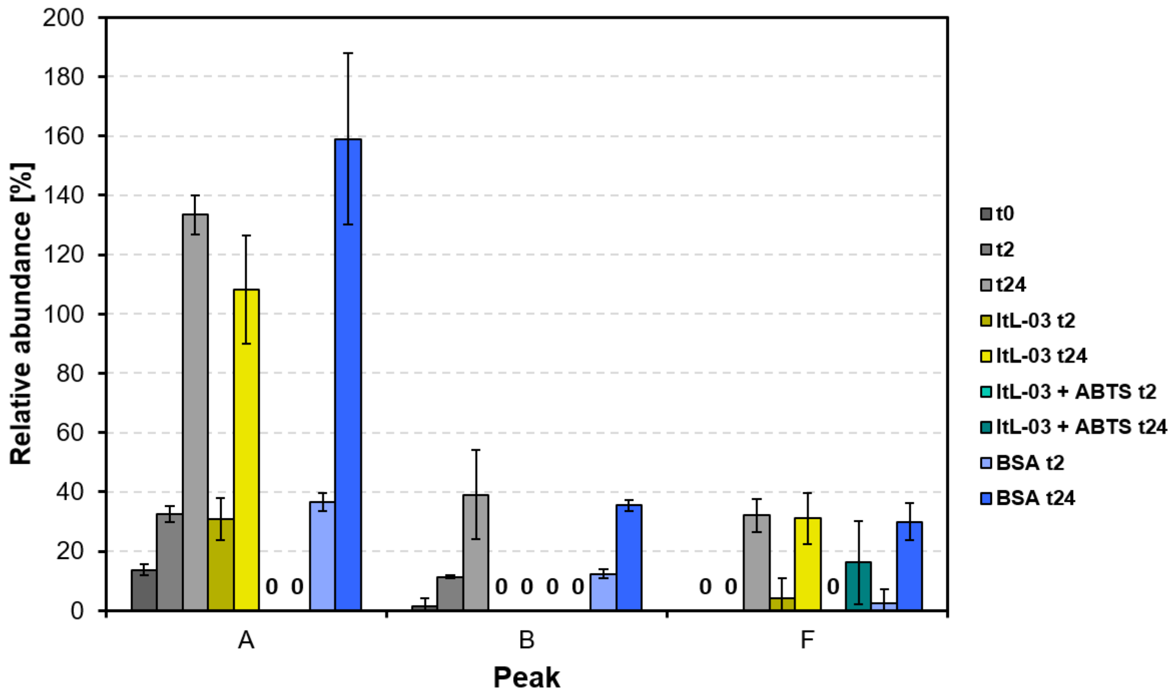


**Figure S18.** The relative abundance of peaks A, B, and F from the same experiment in 'Fig. S17' is shown. The spectra were normalized using the peak at 7.9 min as a reference. $t_0$, $t_2$, and $t_{24}$ represent controls without enzyme at the beginning, between, and end of the incubation period. Peaks A and B were observed in the BSA and control samples; however, peak A was absent in the ItL-03/ABTS samples, and peak B was additionally absent in all ItL-03 samples. Peak F was detected in most samples after 24 hours. Error bars represent the standard deviation ($n$ = 3).

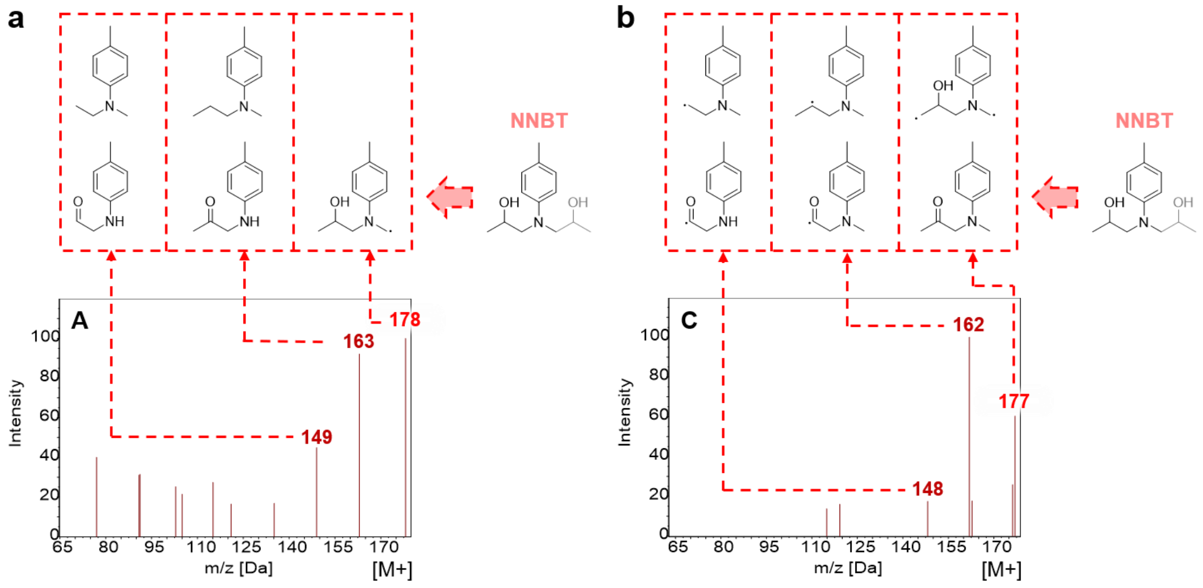


**Figure S19.** Proposed species derived from RTM6 epoxy that may be degraded by ItL-03 are based on GC-MS chromatogram shown in Fig. S17, specifically for the peaks corresponding to (a) A (12.4 min). and (b) C (13.5 min). These ions may originate from NNBT, and the *m/z* pairs of 177 and 178 likely result from the protonation and deprotonation of the same species, as they exhibit similar ionization patterns.

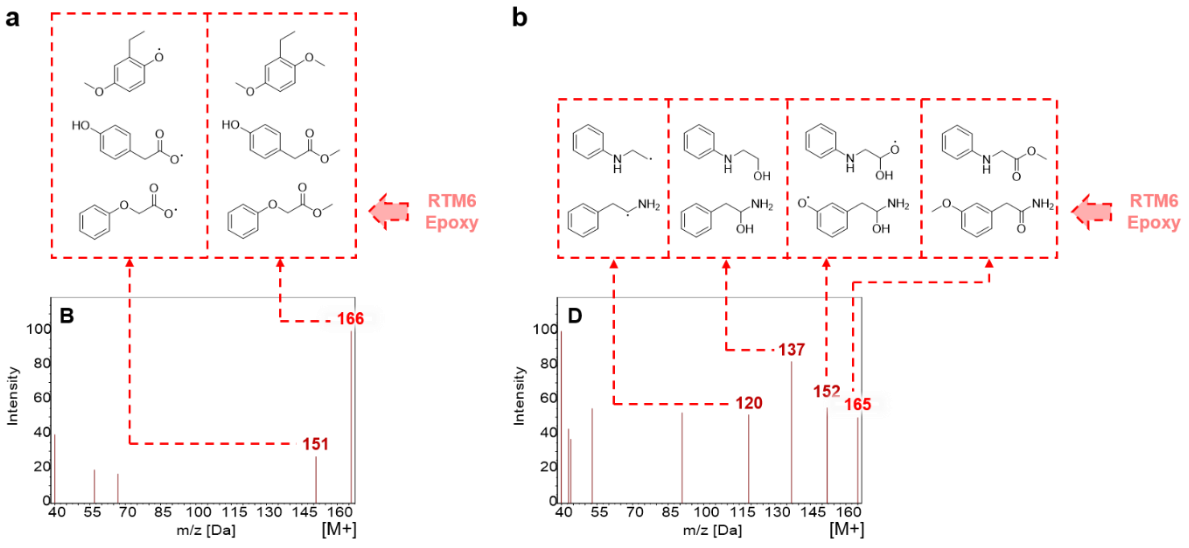


**Figure S20.** Proposed species derived from RTM6 epoxy that may be degraded by ItL-03 are based on GC-MS chromatogram shown in Fig. S17, specifically for the peaks corresponding to (a) B (13.2 min) and (b) D (10.1 min). These ions may originate from the amine epoxy resins that are decomposed by either pre-treatment or the enzyme. The *m/z* pairs of 165-166 could result from the protonation and deprotonation of the same species, as they exhibit similar ionization patterns, or they could represent completely different species.

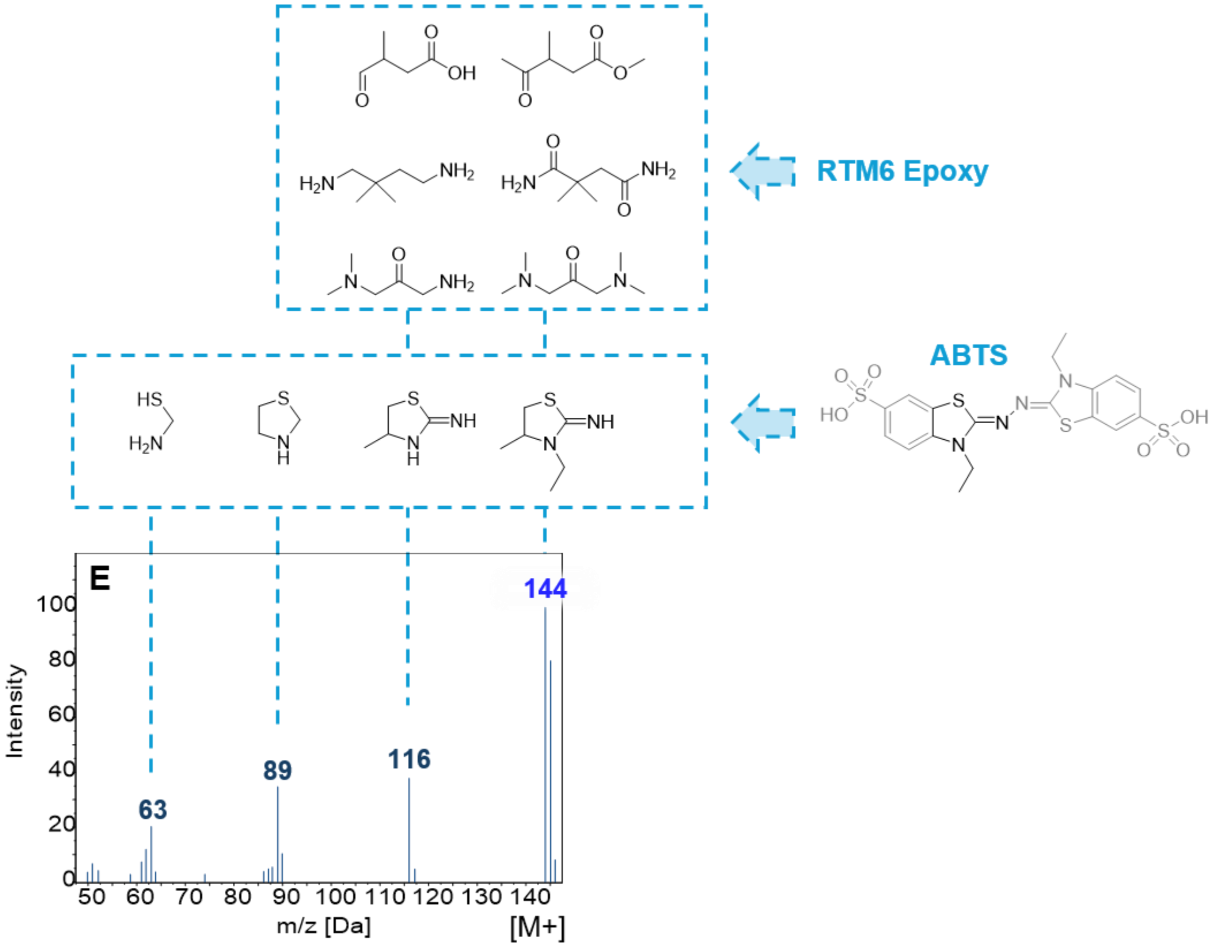


**Figure S21.** Proposed species derived from RTM6 epoxy that may be degraded by ItL-03 in combination with ABTS are based on GC-MS chromatogram shown in Fig. S17, specifically for peak E (14 min). This compound is most likely derived from the fraction of ABTS, as it exhibits ionization patterns consistent with the ABTS mass fraction. Another possibility is that it results from the decomposition of epoxy resins.

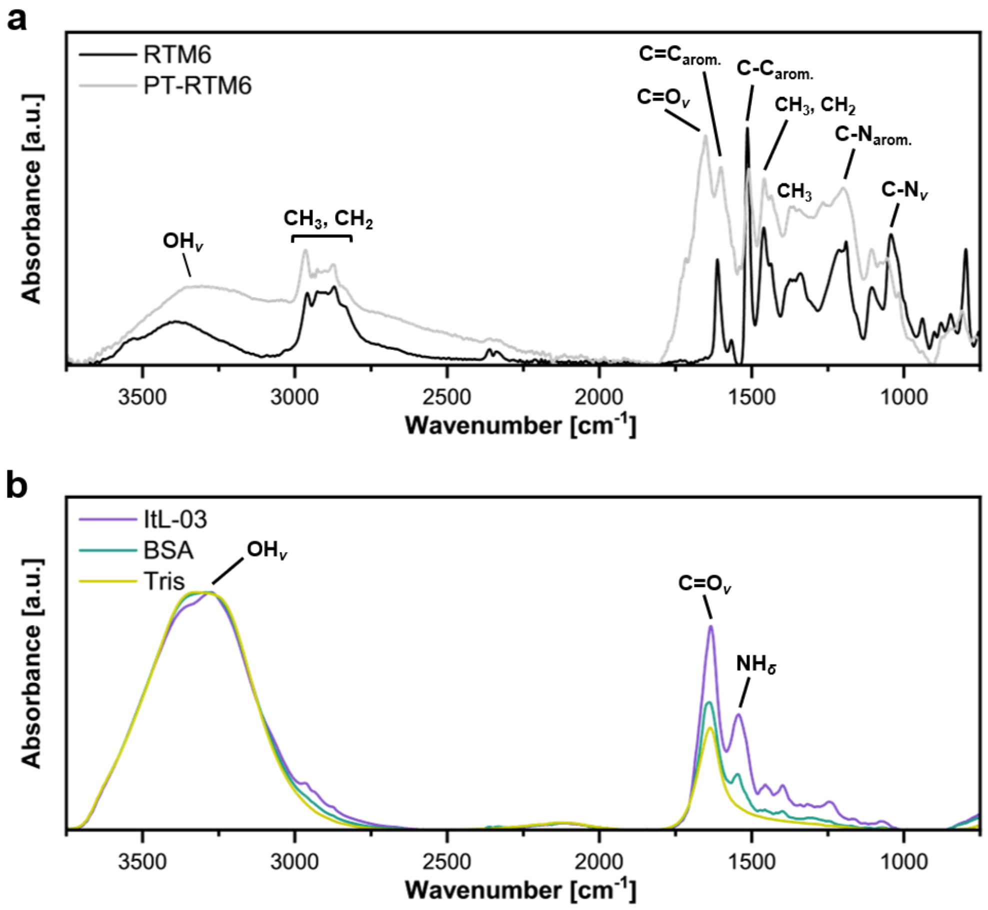


**Figure S22.** Controls for FTIR analysis. (a) Non-treated epoxy powder (RTM6) compared to pre-treated epoxy (PT-RTM6). (b) laccase ItL-03, BSA, and tris buffer that was used for the enzyme stock solution. The corresponding functional groups are indicated. *ν* — stretching; *δ* — deformation.

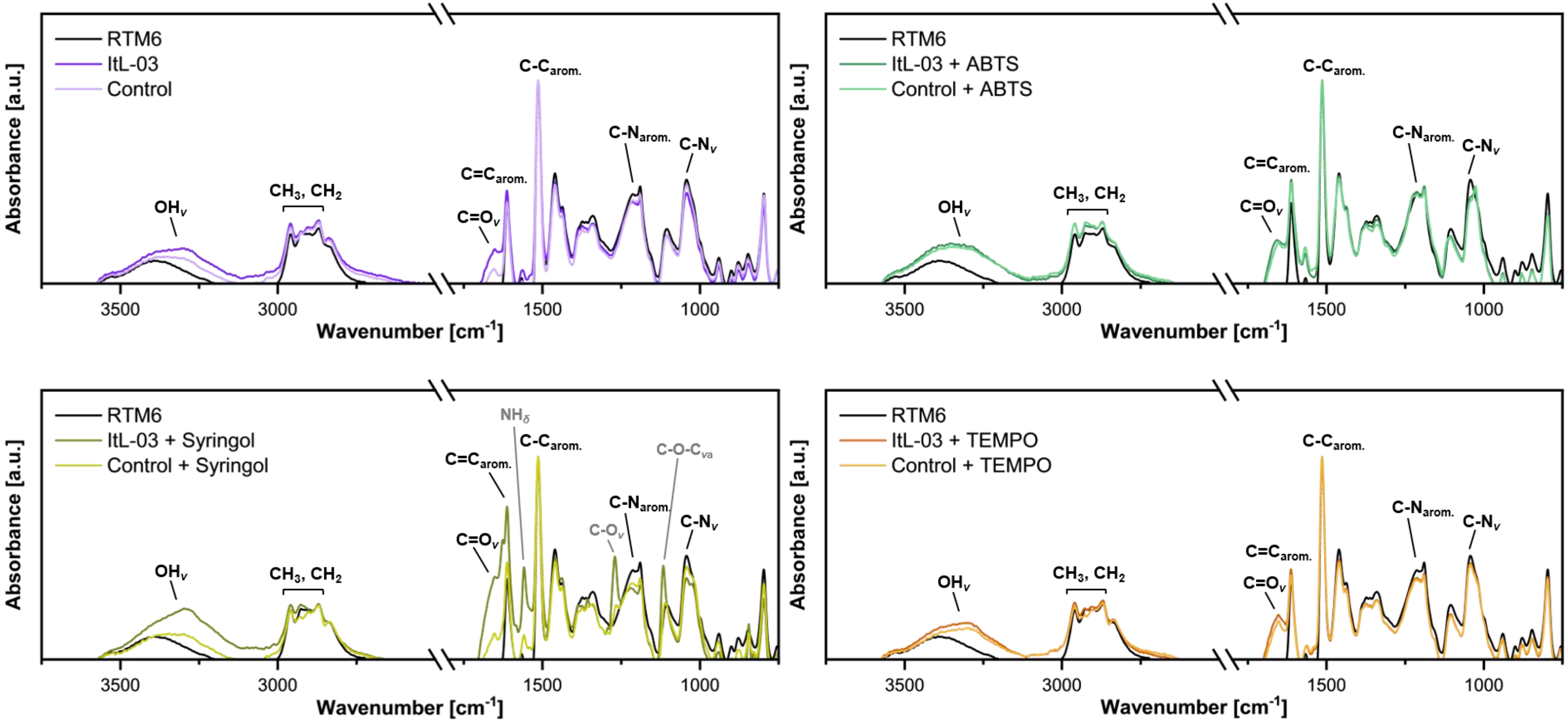


**Figure S23.** FTIR analysis of the degradation activity of ItL-03 after 5 days on non-treated epoxy resins (RTM6) with and without mediators: ABTS, syringol, and TEMPO. BSA was used as a control. The corresponding functional groups are indicated. *v* — stretching; *v*a — asymmetric stretching; *δ* — deformation. Most ItL-03 exhibits activity similar to that of the control, suggesting that the laccase barely degrades the epoxy, even in the presence of mediators. The exception is that ItL-03/syringol shows slightly more activity than the control, particularly in relation to the functional groups C-$N_{arom.}$ and C-$N_{v}$.

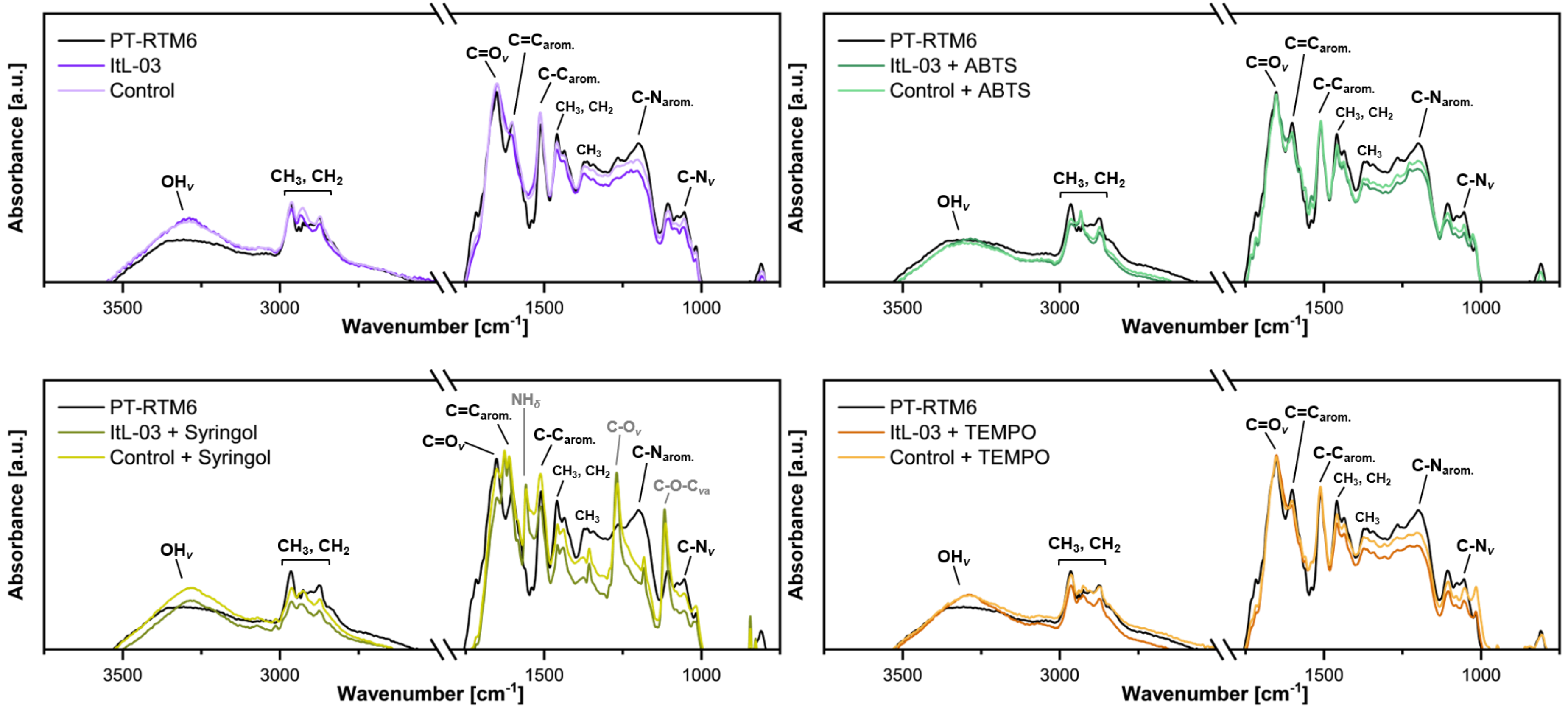


**Figure S24.** FTIR analysis of the degradation activity of ItL-03 after 5 days on pre-treated epoxy powder (PT-RTM6) with and without mediators: ABTS, syringol, and TEMPO. BSA was used as a control. The corresponding functional groups are indicated. *v* — stretching; *v*a — asymmetric stretching; *δ* — deformation. ItL-03, with or without the mediators, can slightly modify the pre-treated epoxy, which shows changes compared to the control, particularly concerning the functional groups C-$N_{arom.}$ and C-$N_v$.
